# Tautomer-specific deacylation and Ω-loop flexibility explain carbapenem-hydrolyzing, broad-spectrum activity of the KPC-2 β-lactamase

**DOI:** 10.1101/2022.11.11.516172

**Authors:** Catherine L. Tooke, Philip Hinchliffe, Michael Beer, Kirill Zinovjev, Charlotte K. Colenso, Christopher J. Schofield, Adrian J. Mulholland, James Spencer

**Affiliations:** School of Cellular and Molecular Medicine, Biomedical Sciences Building, University Walk, University of Bristol, Bristol, BS8 1TD, United Kingdom; Centre for Computational Chemistry, School of Chemistry, Cantock’s Close, University of Bristol, Bristol, BS8 1TS, United Kingdom; School of Biochemistry, Biomedical Sciences Building, University Walk, University of Bristol, Bristol, BS8 1TD, United Kingdom; Departamento de Química Física, Universitat de València, Burjassot, 46100, Comunitat Valenciana, Spain; Chemistry Research Laboratory, Department of Chemistry and the Ineos Oxford Institute for Antimicrobial Research, Mansfield Road, University of Oxford, Oxford, OX1 3TA United Kingdom

## Abstract

KPC-2 (*Klebsiella pneumoniae* carbapenemase-2) is a globally disseminated serine-β-lactamase (SBL) responsible for extensive β-lactam antibiotic resistance in Gram-negative pathogens. SBLs inactivate β-lactams via a mechanism involving a hydrolytically labile covalent acyl-enzyme intermediate. Carbapenems, the most potent β-lactams, evade activity of many SBLs by forming long-lived inhibitory acyl-enzymes; however, carbapenemases such as KPC-2 efficiently catalyze deacylation of carbapenem-derived acyl-enzymes. We present high-resolution (1.25-1.4 Å) crystal structures of KPC-2 acyl-enzymes with representative penicillins (ampicillin), cephalosporins (cefalothin) and carbapenems (imipenem, meropenem and ertapenem), obtained utilizing an isosteric deacylation-deficient mutant (E166Q). Mobility of the Ω-loop (residues 165–170) negatively correlates with antibiotic turnover rates (*k*_cat_), highlighting the role of this region in positioning catalytic residues for efficient hydrolysis of different β-lactams. Carbapenem-derived acyl-enzyme structures reveal predominance of the Δ1-(2*R*) imine tautomer, except for the imipenem acyl-enzyme, which is present in dual occupancy in both Δ1-(2*R*) and (2*S*) configurations. Quantum mechanics/molecular mechanics (QM/MM) molecular dynamics simulations of deacylation of the KPC-2:meropenem acyl-enzyme, using an adaptive string method (ASM), show that the Δ1-(2*R*) isomer has a 7 kcal/mol higher barrier for the (rate-determining) formation of the tetrahedral deacylation intermediate than the Δ2 tautomer. The simulations identify tautomer-specific differences in hydrogen bonding networks involving the carbapenem C-3 carboxylate and the deacylating water, that, together with stabilization by protonated N-4 of accumulating negative charge during oxyanion formation, accelerate deacylation of the Δ2-enamine acyl-enzyme compared to the Δ1-imine. Taken together, our data show how the flexible Ω-loop helps confer broad spectrum activity upon KPC-2, while carbapenemase activity stems from efficient deacylation of the Δ2-enamine acyl-enzyme tautomer. Differentiation of the barriers associated with deacylation of these subtly different β-lactam isomers further identifies ASM as a sensitive method for calculation of reaction energetics that can accurately model turnover and, potentially, predict the impact of substrate modifications or point mutations upon activity.

## Introduction

β-Lactams are the most prescribed antibiotics worldwide for the treatment of serious healthcare-associated infections (HAIs) caused by Gram-negative bacterial pathogens.^1-2^ Many aspects of healthcare are threatened by increasing resistance to these antibiotics. Around 2.8 million antibiotic-resistant infections occur each year in the USA^3^ and 1.27 million deaths worldwide are attributed to bacterial resistance to antibiotics, and these numbers are growing.^4^ The most important mechanism of β-lactam resistance in Gram-negative bacteria is the production and activity of serine β-lactamases (SBLs), a diverse class of hydrolytic enzymes which inactivate all classes of β-lactam antibiotics.^5^ SBLs are split into three Ambler classes, A, C and D, based on sequence and mechanism. In all classes, SBL catalyzed β-lactam hydrolysis proceeds through two main steps: attack of the nucleophilic serine on the β-lactam carbonyl carbon to form a covalent acyl-enzyme intermediate with cleavage of the β-lactam amide bond (acylation),^6-7^ followed by deacylation through attack of an activated water molecule on the acyl-enzyme complex carbonyl to release the inactivated hydrolyzed antibiotic^8^ (**Figure 1**).

**Figure 1.**
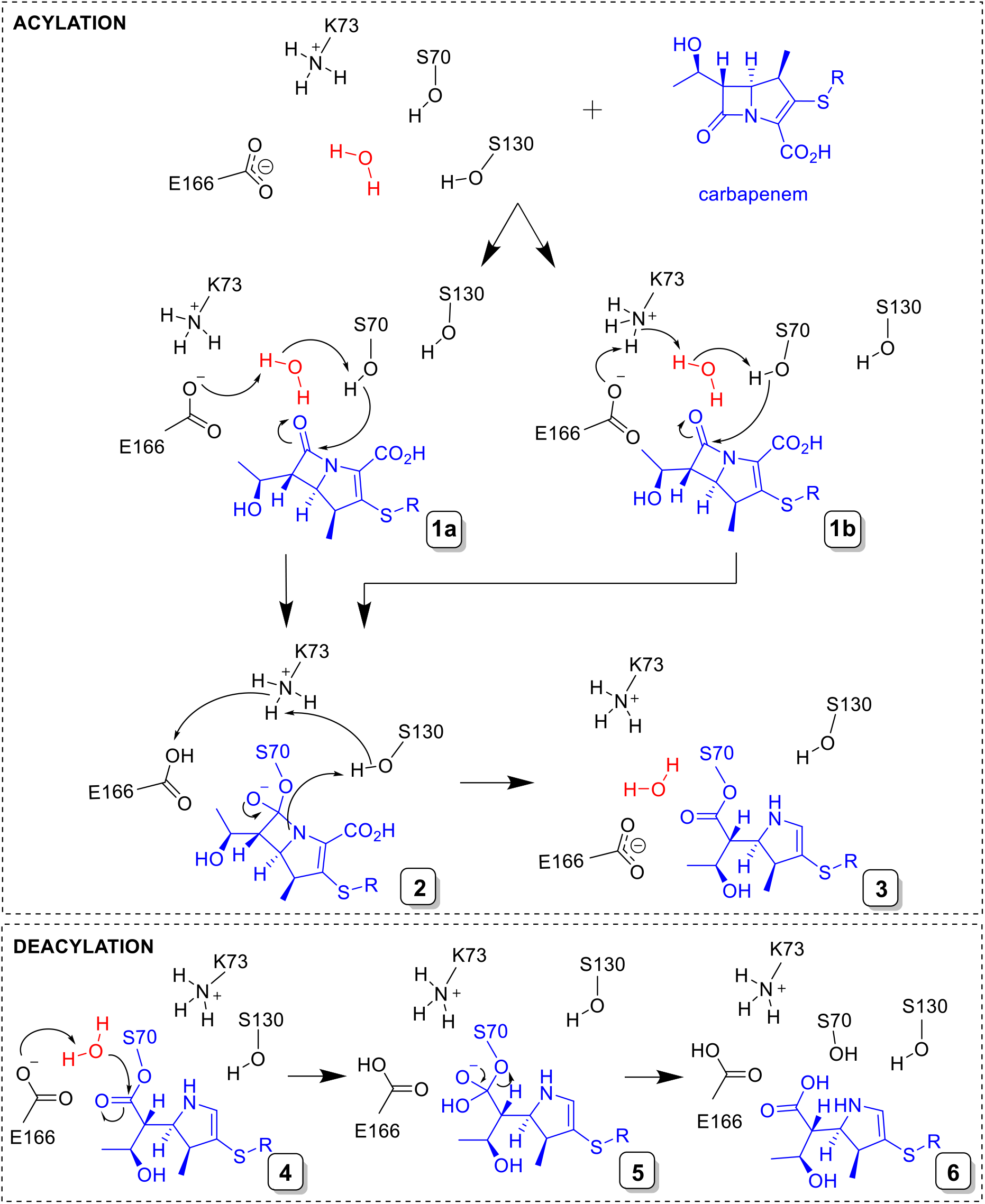
Reaction of carbapenems with class A serine β-lactamases. *Top*, acylation; binding of a 1β-methyl carbapenem (blue) to the class A SBL active site is followed by acylation^6^; Ser70 is activated via an active site water (red) with either Glu166^7^ (1a) and/or Lys73 (1b) acting as a general base, and subsequent nucleophilic attack on the C-7 carbonyl forms a tetrahedral intermediate (2) which reacts to give the covalent acyl-enzyme complex (3, for clarity only the Δ2-enamine is shown). **B**. *Bottom*, deacylation^8^; proton transfer from the deacylating water molecule (red) to Glu166 enables nucleophilic attack on the C-7 carbonyl of the acyl-enzyme complex (4) resulting in a second tetrahedral intermediate (5) that collapses to give the hydrolyzed product (6).

Carbapenems are the most potent β-lactams for the treatment of severe HAIs caused by opportunistic Gram-negative species. Carbapenems contain a five-membered pyrroline ring fused to a four-membered β-lactam ring and were first discovered as natural products, such as thienamycin, but those used clinically are produced by total synthesis.^9^ Their pyrroline ring differentiates them from the penicillins and cephalosporins, which contain β-lactam rings fused to thiazolidine and dihydrothiazine rings, respectively. Furthermore, the 6α-hydroxyethyl carbapenem substituent, which differs from the typically larger β-substituent groups attached to C-6/C-7 in penicillin and cephalosporin antibiotics (**Figure 2**), is thought to protect carbapenems against hydrolysis by most β-lactamases.^9^ Indeed, carbapenems were originally discovered as both antimicrobials and β-lactamase inhibitors, and are known to resist deacylation by many SBLs (e.g. the class A TEM, SHV and CTX-M enzymes), by forming long-lived, acyl-enzyme complexes.^5^ The development of carbapenems to improve pharmacological activity and potency has involved addition of a 1β-methyl group to protect against hydrolysis by human dehydropeptidase-I^9^ and alternative substituents at the (*sp*^2^ hybridized) C-2 position (R1 groups, **Figure 3**).

**Figure 2.**
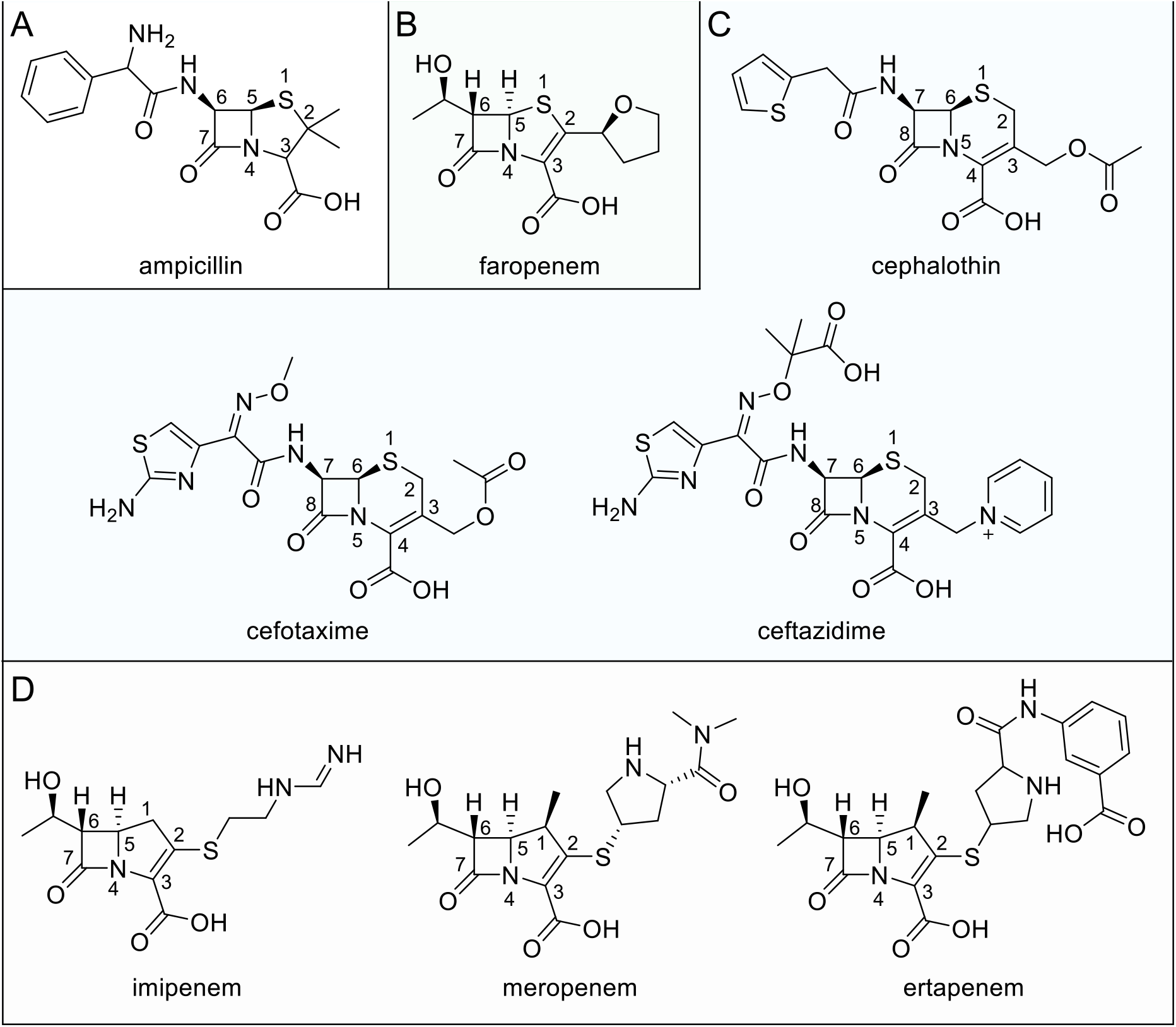
Structures of β-lactam antibiotics used in this study. **(A)** Ampicillin (penicillin/penam). **(B)** Faropenem (penem). **(C)** Cephalothin, cefotaxime and ceftazidime (cephalosporins). **(D)** Imipenem, meropenem and ertapenem (carbapenems).

**Figure 3.**
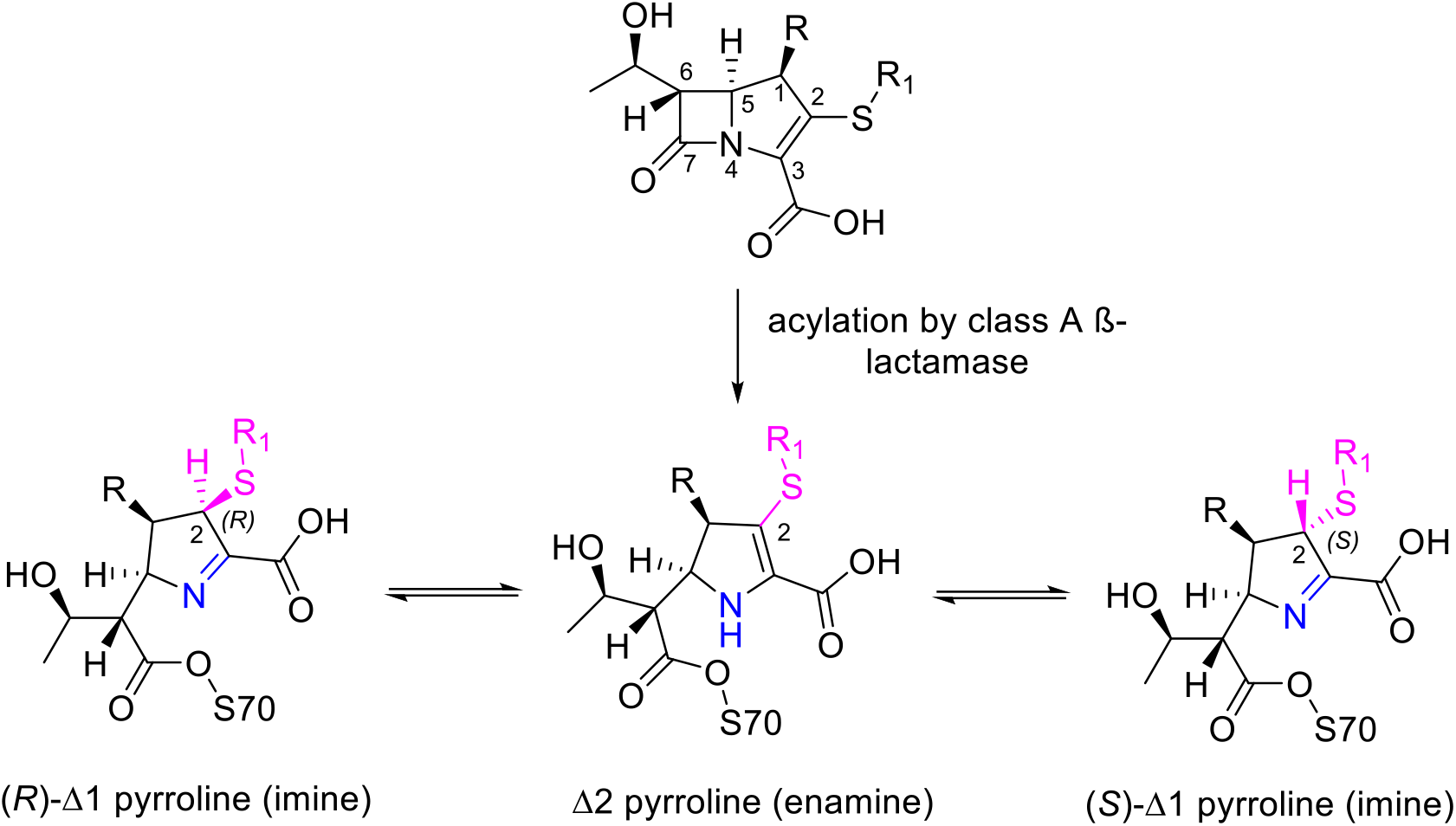
Carbapenem tautomerization in acyl-enzyme complexes of class A β-lactamases.

The presence of a double bond in the carbapenem pyrroline ring permits tautomerization of the acyl-enzyme complex formed between Δ1-pyrroline and Δ2-pyrroline forms (**Figure 3**). Migration of the double bond from C-2=C-3 (Δ2, enamine) to C-3=N (Δ1, imine), with associated protonation at C-2, gives rise to the possibility of alternative enamine and imine hydrolysis products, with the latter in either the 2*R*-or 2*S*-configurations. For example, in the class A SBL SHV-1, which lacks carbapenemase activity, Raman spectroscopy has revealed rapid deacylation to give a Δ2 enamine, and prolonged stability of Δ1, leading to the suggestion that accumulation of the latter is the basis for poor turnover.^10^ In this and similar enzymes deacylation of carbapenem acyl-enzymes is believed to be retarded by interaction of the 6α-hydroxyethyl group with the deacylating water molecule, so reducing its nucleophilicity.^11^

Since the introduction of carbapenems in the early 1980s, β-lactamases that efficiently inactivate these antibiotics (carbapenemases) have emerged. These include carbapenem-hydrolyzing SBLs that now significantly challenge the clinical efficacy of all members of this potent β-lactam antibiotic class.^12^ Such resistance is now common with carbapenem-resistant *Acinetobacter baumannii, Pseudomonas aeruginosa* and *Enterobacterales* all now listed as WHO priority pathogens.^13-15^ One such carbapenem-hydrolyzing enzyme, *Klebsiella pneumoniae* carbapenemase-2 (KPC-2), is a class A SBL first identified from a carbapenem-resistant *Klebsiella pneumoniae* isolate from North Carolina.^13^ KPC-2 is now present worldwide in numerous pathogenic Gram-negative organisms.^16^ KPC-2 is described as a ‘versatile β-lactamase’ because it efficiently hydrolyzes most penicillins, cephalosporins, carbapenems, and resists the action of inhibitors based on the β-lactam scaffold.^5, 9, 12^

In KPC-2, and other class A SBLs, the active site (containing the nucleophilic Ser70) is bordered by the flexible Ω-loop, where the general base for deacylation (Glu166) resides.^5, 7, 11^ In crystal structures of unliganded KPC-2, Ser70 and Glu166 participate in conserved H-bond networks that also involve Lys73, Ser130, Asn170 and an active site water molecule apparently positioned for the deacylation reaction (**Figures 1 and S1**). We recently showed how plasticity in and around the KPC-2 active site enables the accommodation of more ‘bulky’ antibiotics such as the expanded-spectrum oxyimino-cephalosporin ceftazidime, although this adversely affects turnover rates due to displacement of the Ω-loop and consequent imperfect repositioning of Glu166.^11^

Quantum mechanics/molecular mechanics (QM/MM) simulations of deacylation can be used as a “computational assay” to assess the ability of class A β-lactamases to hydrolyze carbapenems and to discriminate between carbapenemases and non-carbapenemases on the basis of calculated reaction barriers.^17^ However, while these and other computational approaches can identify some determinants of carbapenem turnover (for example, in carbapenem-hydrolyzing enzymes, interaction with Asn132 is proposed to position the 6α- hydroxyethyl substituent away from direct interaction with the deacylating water^18,19^) the availability of experimental information, in particular high-resolution structures of acyl-enzyme complexes, remains limited. As a consequence, a comprehensive description of the basis for the different activities of class A β-lactamases towards carbapenems remains to be achieved. Indeed, the basis for the carbapenem-hydrolyzing activity of KPC and related enzymes, which distinguishes them from the great majority of class A β-lactamases, and in particular how the KPC-2 active site ensures efficient carbapenem deacylation, is poorly understood.^11^

Here, we present crystal structures of carbapenem acyl-enzyme complexes of the deacylation-deficient KPC-2 Glu166Gln mutant (KPC-2^E166Q^), which unexpectedly are predominantly present as the Δ1, rather than Δ2, tautomer. Based upon this information, we explored the dynamics of the acyl-enzyme complex formed by the carbapenem meropenem with wild-type KPC-2 using 1.5 μs of molecular mechanics molecular dynamics (MM MD) simulations, and apply QM/MM calculations to investigate deacylation of both the meropenem Δ1 and Δ2 tautomers using the adaptive string method.^20^ The results indicate that a key factor underlying the efficient hydrolysis of multiple β-lactam classes by KPC-2 is the ability of the Ω-loop to enable efficient positioning of the deacylating water (DW) in the respective acyl-enzymes. Furthermore, efficient deacylation of carbapenem-derived acyl-enzymes in the Δ2 enamine tautomer is enabled by hydrogen bonding networks involving the C-3 carboxylate and 6α-hydroxyethyl groups and DW, and contrasts with much weaker activity towards the Δ1 configuration. These data identify the basis for the broad spectrum of activity of KPC-2 and how this enzyme, and by implication other class A carbapenemases, efficiently deacylates carbapenem acyl-enzymes. Our results will inform the design of both new generations of β- lactamase inhibitors and of new β-lactams able to evade the versatile, broad-spectrum activity of KPC-2.

## Methods

### Enzyme assays

All enzyme assays were followed at 25 °C in 10 mM HEPES pH 7.5, 150 mM NaCl in Greiner half area 96-well plates and a Tecan Infinite® 200 pro microplate reader. Steady-state kinetic parameters were calculated by measuring β-lactam antibiotic hydrolysis (ampicillin Δ235 ε =-900, cefalothin Δ262 ε=-7660, imipenem Δ299 ε=-9000, meropenem Δ297 ε=-6500 and ertapenem Δ295 ε=-7112) with 10 nM KPC-2 with 50 ug/mL BSA. Initial rates (*V*_0_) of β-lactam hydrolysis were plotted against the concentration of antibiotic and kinetic parameters were calculated and analyzed by least-squares fitting to the Michaelis-Menten equation in GraphPad Prism 6, (GraphPad Software, La Jolla California USA, www.graphpad.com).

### Protein crystallization, antibiotic complex generation and X-ray diffraction data collection

KPC-2 and KPC-2^E166Q^ were produced in recombinant *E. coli* using the pET28a T7 expression vector and were purified and crystallized as described previously.^11^ KPC-2^E166Q^ crystals were soaked with solutions of 10 mM ampicillin (Sigma), 15 mM cefalothin (Sigma), 15 mM imipenem (Sigma), 30 mM meropenem (Sigma) and 30 mM ertapenem (MedChemExpress, USA) dissolved in mother liquor (2.0 M ammonium sulphate, 5% ethanol) supplemented with 20-30% glycerol. Crystals were soaked from 5 minutes to several hours. The datasets presented here were of the best quality of those obtained (determined by resolution, ligand occupancy and RSCC, alongside overall data collection and refinement statistics) and were collected after soaking crystals for 2.5 hours (KPC-2^E166Q^:ampicillin and KPC-2^E166Q^:cefalothin, KPC-2^E166Q^:imipenem and KPC2^E166Q^:meropenem) and 3 hours (KPC2^E166Q^:ertapenem). Diffraction data were collected at the ALBA synchrotron beamline BL13 XALOC. Images were indexed and integrated using XDS and subsequently scaled in AIMLESS (CCP4 suite).^21^ Crystallographic phases were calculated in Phaser^21-22^ using PDB 6Z21 (crystal structure of apo KPC-2^E166Q^) as a molecular replacement solution. Initial refinements in REFMAC5^23^ confirmed *F*_o_-*F*_c_ electron density consistent with a single ligand bound at the active site, prior to further rounds of refinement in Phenix.refine^24^ and manual model building in Coot.^21, 25^ Geometry restrains for antibiotic-derived ligands were calculated using eLBOW in Phenix^24^ and omit maps were generated in Phenix^24^ from the final model in the absence of the antibiotic. Ligand occupancies were initially manually assigned based upon visual inspection of electron density and subsequently refined in Phenix^24^ with at least 10 rounds of refinement. Figures were generated in Pymol.^26^

### Molecular dynamics simulations

Crystal structures of KPC-2 (PDB 5UL8^27^), KPC-2^E166Q^ (PDB 6Z21^11^) and KPC-2^E166Q^:meropenem were used as starting structures for molecular simulations. For simulations of the wild-type KPC-2, Q166 was edited to E166^11^ in Coot.^25, 28^ The Δ2 meropenem-derived acyl-enzyme complex was modelled in Coot into KPC-2 using the electron density of the Δ1-(2*R*) meropenem-derived acyl-enzyme complex as a guide. The Δ1-(2*R*) complex of KPC-2:meropenem was deemed to be the most representative carbapenem complex structure for simulation as all meropenem atoms were well defined by the experimental electron density and meropenem has both a smaller C-2 substituent than ertapenem and contains a C-1 methyl group (absent from imipenem). In addition, the Δ1-(2*R*) complex was the most frequently observed carbapenem tautomer and consistently refined with the highest occupancy. The resulting protein coordinates (KPC-2^E166Q^, KPC-2, KPC-2^E166Q^:Δ1-meropenem and KPC-2:Δ1-meropenem, KPC-2^E166Q^:Δ2-meropenem and KPC-2:Δ2-meropenem) were parameterized for molecular simulations. All crystallographically observed water molecules were included in the structures. The protonation states of titratable residues were determined using the PropKa 3.1 server.^29^ Hydrogens were added in tleap (AMBER16^30^) and the systems were solvated using a 10 Å waterbox (TIP3P) with overall charges neutralized by addition of Na^+^ or Cl^−^ ions replacing bulk water molecules. Atomic charges for the meropenem (Δ1/Δ2) acyl-enzymes were generated using restrained electrostatic potential (RESP) fitting as implemented in the RED server.^31^ All structures underwent a standard energy minimization (600 steps of steepest descent and 600 steps of conjugate gradient), heating (25 K to 298 K in 20 ps) and equilibration by MM MD (1 ns) protocol. The structures were then simulated using MM MD in the AMBER16 simulation package using the ff14SB MM forcefield for protein, the TIP3P-Ew water model and the general AMBER force field (GAFF). All 6 protein systems were simulated in triplicate for 100 ns; KPC-2, KPC-2:Δ1-meropenem and KPC-2:Δ2-meropenem were further simulated in triplicate runs of 500 ns (1.5 μs in total). RMSD, RMSF, clustering, distance and dihedral analyses were performed in CPPTRAJ in AMBER16. RMSD calculations were performed using the first frame (1 ps) as the reference.

### QM/MM calculations of carbapenem deacylation (tetrahedral intermediate formation)

An implementation of the adaptive string method (ASM) with AMBER18^32^ was used to calculate the minimum free energy path (MFEP).^20^ Two collective variables were chosen to monitor reaction progress: the distance between the transferred proton of the DW and the Glu166 side chain (collective variable 1: rx = d(OεGlu166–HDW) and the distance between the oxygen of the DW and meropenem C-7 carbon (collective variable 2: ry = d(C-7meropenem–ODW)) (**Figure S2)**. The ASM applies an on-the-fly string method for defining the position of the minimum free energy pathway (MFEP). The position of the string nodes (after sufficient string position optimization) is then used to define the reaction coordinate (RC) and umbrella sampling (US) windows, used to calculate the potential of mean force (PMF) of the proposed reaction pathway. The ASM differs from conventional umbrella sampling (US) as it does not use user-specified RC and US windows, providing a flexible description of the reaction.^20^ For simplification, we refer to this method as ASM and conventional umbrella sampling as US, despite both methods employing umbrella sampling to obtain data for PMF calculation.

Acyl-enzyme complex and tetrahedral intermediate structures for string method calculations were established through US calculations of the minimum free energy path (MFEP) using the same reaction coordinates (**Supporting note 1**). The QM calculations used the semi-empirical SCC-DFTB2 level of theory, with the ff14SB MM forcefield for the protein, the TIP3P-Ew for the DW water model, and the General AMBER Force Field (GAFF) for those parts of meropenem not included in the QM region. The QM region (modelled with an overall charge of –2) comprised the deacylating water (DW), the side chains of Ser70 and Glu166, and the carbapenem scaffold (**Figure S2**). The string was composed of 28 nodes. Each initial starting structure (acyl-enzyme complex) was obtained from 300 ps of QM/MM MD, and end structure (tetrahedral intermediate) by 20 ps sampling in an umbrella sampling simulation of the at the desired reaction coordinate. No nodes were fixed to allow the system to completely relax, and no additional restraints were placed on any atoms.

During the string optimization phase of the simulations, string convergence was monitored by calculating the RMSD of the string at each step compared to all the previous states of the string. In all cases convergence of the string was reached by 50 ps and converged parameters for the string were obtained by averaging over 10,000 steps from when the string was judged to have reached convergence.

60 ps of umbrella sampling was then run with these parameters and the potential of mean force (PMF) calculated with umbrella integration.^33^ This procedure was repeated three times for the Δ1 and Δ2 tautomers, using a different initial and final string node (obtained from independent simulations) in each repeat. The MFEP was analyzed over a set of 28 trajectories, one for each node. TS trajectories were formed from frames with a PMF within 0.05 kcal/mol of the calculated barrier. Each trajectory was analyzed using CPPTRAJ from AmberTools16.^34^

## Results

### Kinetic and structural analysis of antibiotic hydrolysis by KPC-2

KPC-2 is a carbapenemase that also hydrolyzes a wide variety of other β-lactams, including penicillins and cephalosporins, as demonstrated by steady-state kinetic data (**Table 1** and previous work^35-38^). KPC-2 efficiently catalyzes the hydrolysis of the three carbapenems tested here, although the *K*_M_ value for the hydrolysis of imipenem (which contains a 1β-hydrogen rather than 1β-methyl group) is an order of magnitude larger those for the 1β-methyl-substituted carbapenems, ertapenem and meropenem (*K*_M_ values 72, 7.1, and 8.6 μM, respectively). The *K*_M_ values also increase for β-lactam antibiotics, that is cephalosporins and penicillins, with larger C-6/C-7 groups in the β-orientation, compared to the relatively small 6-hydroxyethyl group in the α-orientation that is present in carbapenems and penems (**Table 1)**.

**Table 1.**
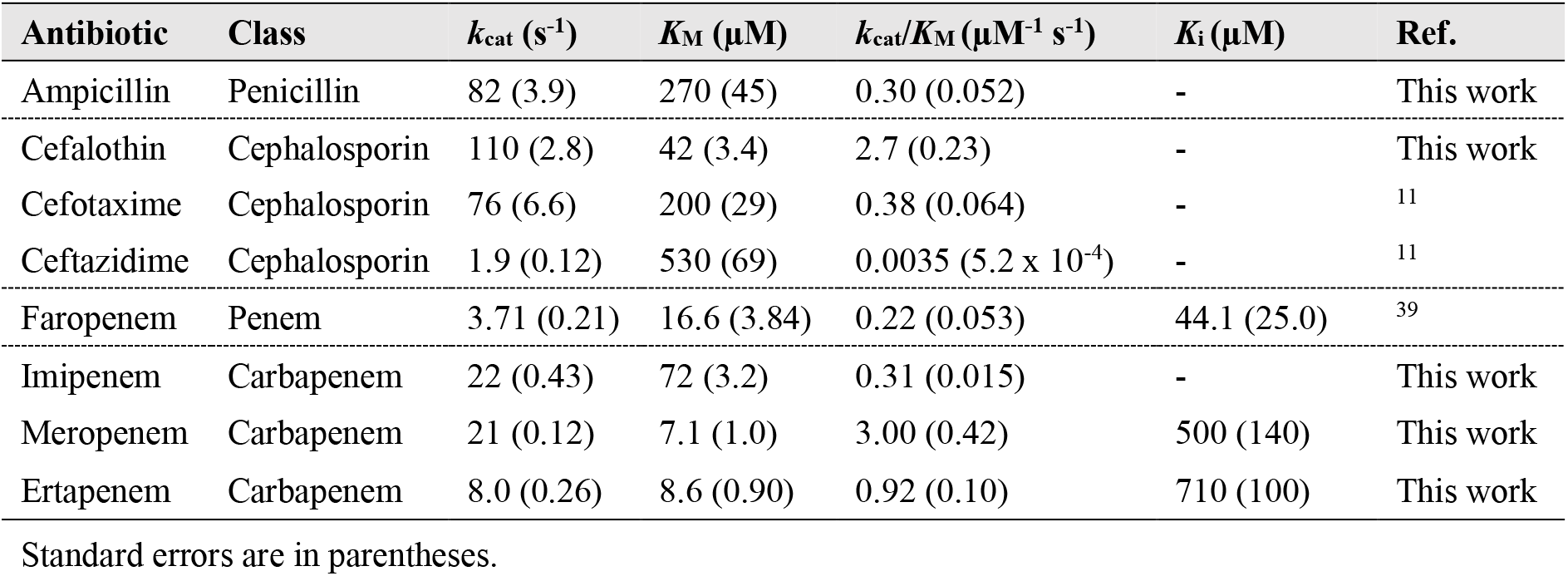
KPC-2 steady-state kinetics.

To investigate the interactions of members of these different antibiotic classes with the KPC-2 active site we have utilized the isosteric Glu166Gln substitution (KPC-2^E166Q^)^11^ to slow deacylation of the acyl-enzyme (allowing resolution of the acyl-enzyme intermediate). Preformed KPC-2^E166Q^ crystals were soaked with solutions of ampicillin, cefalothin, imipenem, meropenem or ertapenem, representing three different β-lactam antibiotic classes (penicillins, cephalosporins and carbapenems) for a range of exposure times, and diffraction data were collected. The resulting data extended to high resolution (1.25 - 1.4 Å) (**Table S1**), with clear *F*_o_ - *F*_c_ difference density in the KPC-2 active sites enabling confident modelling of the antibiotic-derived complexes covalently linked with the nucleophilic Ser70 (**Figure 4**). Real-space correlation coefficient (RSCC) values of 0.89-0.97 (as calculated by the wwPDB validation server^40^, **Table S2**) support reliable fitting of the modelled ligands to the experimental electron density maps. As observed in other crystal structures of SBL:cephalosporin acyl-enzyme complexes,^11^ the 3’
s leaving group of the cefalothin-derived product could not be modelled and has most likely been eliminated during rearrangement of the acyl-enzyme. The cefalothin C-7 thiophenylacetamido substituent was modelled in two ring flipped conformations (occupancies 0.91/0.09) that differ in the orientation of the thiophenyl ring (**Figure S3**). The penicillin C-6 2-amino-2-phenylacetamido substituent was modelled in a single conformation in which all atoms could be resolved. By contrast, the C-2 substituents of the three carbapenem-derived acyl-enzymes are less well defined by the experimental electron density. In particular, the benzoic acid derived atoms of the ertapenem side chain could not be modelled, indicating considerable flexibility of the C-2 groups of carbapenem-derived acyl-enzymes within the KPC-2 active site (**Figure S4**). For all the structures presented herein KPC-2 could be modelled as a continuous chain (residues 25-293) with the respective Ω-loops (containing Gln166) all present in single conformations. This contrasts with our previously published structure of the KPC-2^E166Q^:faropenem-derived acyl-enzyme complex, where 2 conformations of Gln166 were observed, oriented either ‘in’ or ‘out’ of the active site.^39^

**Figure 4.**
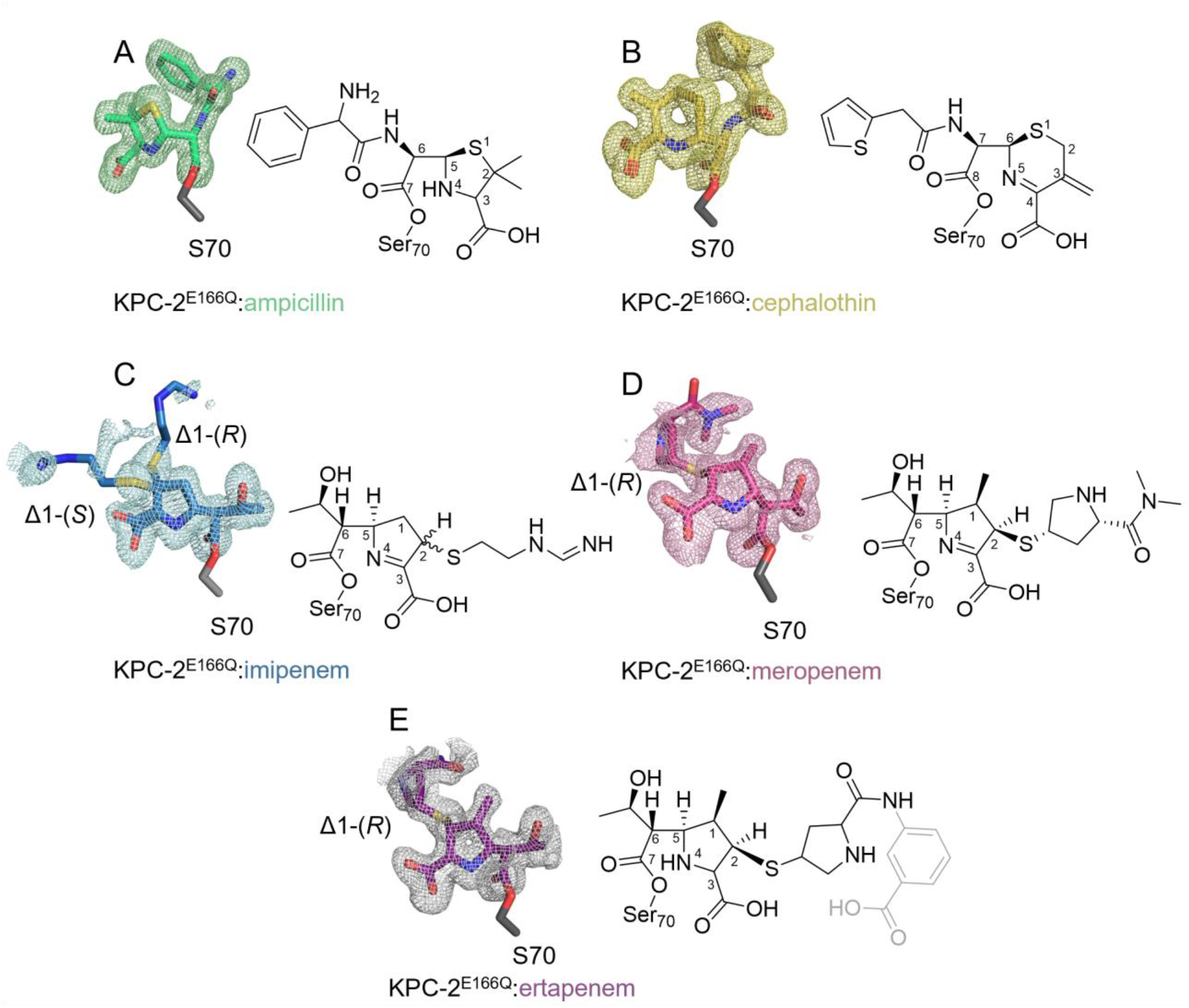
Acyl-enzyme complexes of KPC-2^E166Q^. *F*_o_-*F*_c_ electron density maps (colored mesh, contoured at 3σ) are calculated from the final model with the ligand omitted. **A**. Ampicillin-derived acyl-enzyme (green, PDB:8AKI). **B**. Cefalothin-derived acyl-enzyme (yellow, PDB:8AKJ). **C**. Imipenem-derived acyl-enzyme (blue, PDB:8AKK). Note, dual conformation of the ligand (C-2 atom in both (*R*) and (*S*) configuration) clearly defined by electron density of the R-group sulfur. **D**. Meropenem-derived acyl-enzyme (pink, PDB:8AKL) **E**. Ertapenem-derived acyl-enzyme (purple, PDB:8AKM), atoms in gray were not modelled due to poorly defined electron density in this region.

The antibiotic-derived acyl-enzymes (PDBs 8AKI-8AKM, **Table S1**) are positioned to form hydrogen bonds (H-bonds) with the side chain hydroxyl of the active site residue Ser130, which acts as a H-bond donor to N-4 of bound ampicillin, or as a H-bond acceptor to the imine N-4/N-5 of bound carbapenems and cephalosporins. Similar interactions were also observed in our previous structures of the KPC-2 acyl-enzymes formed on reaction with cefotaxime, ceftazidime^11^ and faropenem^39^. Comparison with the uncomplexed enzyme (PDB: 5UL8^41^), reveals that Asn132 retains the H-bond network with Lys73 and Gln166, while forming an additional H-bond with the C-6/C-7 amide nitrogen of penicillin/cephalosporin, or the 6α-hydroxyethyl hydroxyl in acyl-enzymes of penems/carbapenems.

### KPC-2:carbapenem acyl-enzymes are present in crystals as the Δ1 tautomers

In all cases the carbapenem-derived acyl-enzymes were modeled in the KPC-2 active site as the Δ1-pyrroline (imine) tautomer, as evidenced by the position of the exocyclic sulfur, which at these resolutions/occupancies can be clearly resolved as out of the plane of the pyrroline ring (**Figures 4 and 5**). The C-2 atom is therefore, at least predominantly, *sp*^3^ hybridized in all cases; in the case of imipenem, the carbapenem acyl-enzymes were modeled into the experimental electron density as the (2*R*)*-* and (2*S*)*-*enantiomers in dual occupancy (imipenem occupancy 0.5/0.5 for the (2*R*)- and (2*S*)-forms, respectively) while the 1β-methyl carbapenems meropenem and ertapenem were modeled as the Δ1-(2*R*)-enantiomer only, as in the substrate carbapenem. In all cases, the β-lactam derived C-7 carbonyl of the antibiotic-derived acyl-enzyme is positioned in the oxyanion hole formed by the backbone amides of Ser70 and Thr237, and the 6α-hydroxyethyl oxygen atom is positioned to H-bond with the side chain nitrogen of Asn132 (**Figure 5, Figures S5** and **S6**).

**Figure 5.**
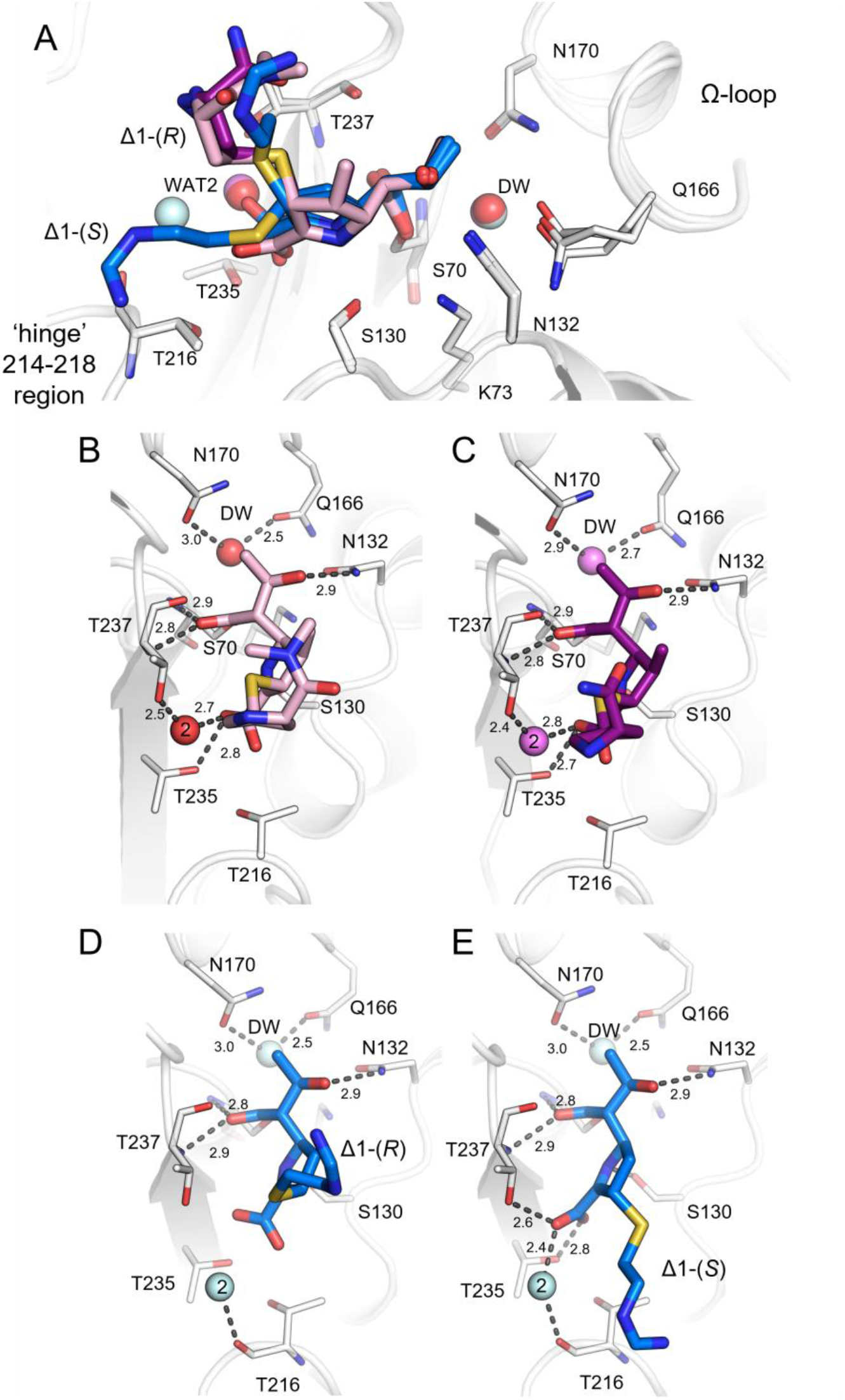
Hydrogen bonding interactions in the KPC-2_E166Q_:carbapenem acyl-enzyme complex structures. **A. S**uperposition of KPC-2^E166Q^ acyl-enzyme complexes with imipenem (blue), meropenem (pink) and ertapenem (purple**) B**. KPC-2^E166Q^:meropenem. **C**. KPC-2^E166Q^:ertapenem acyl-enzyme complexes. **D. and E**. KPC-2^E166Q^:imipenem in the Δ1-(2*R)* and Δ1-(2*S)* configurations. The protein backbone is represented as a white cartoon and active site residues as thin sticks. The deacylating water (DW) and a second active site water interacting with the carbapenem carboxylate (2) are shown as colored spheres. Interactions are shown as dashed grey lines with distances (Å) labeled.

In the meropenem and ertapenem Δ1-(2*R*) acyl-enzymes, the C-3 carboxylate is positioned to interact with the sidechain oxygen atoms of Thr235 and, *via* a water molecule (Wat2, **Figure 5, Figures S5** and **S6**), Thr237. By contrast, in the Δ1-(2*S*)-imipenem acyl-enzyme, the C-3 carboxylate is rotated by 90° with respect to its position in the Δ1-(2*R)-*imipenem acyl-enzyme (**Figure 5**), and makes interactions with Thr237 and Thr235 and, *via* a water molecule (possibly that displaced from the Wat2 position (**Figure 5, Figures S5** and **S6**)) with the backbone carbonyl oxygen of Thr216 in the ‘hinge’ region. The rotation of the C-3 carboxylate group in the Δ1-(2*S*)-imipenem acyl-enzyme is necessary to avoid a steric clash with the exocyclic sulfur of the C-2 substituent. In the Δ1-(2*R*)-imipenem acyl-enzyme, the C-3 carboxylate is in a similar position to that observed in the Δ1-(2*R*) meropenem and ertapenem acyl-enzymes, but the H-bond interactions with the Thr237 and Thr235 side chains and Wat2 are lost (**Figures S5** and **S6**), probably due to the displacement of Wat2, which is necessary to support exchange of Δ1-(2*R*) with its (2*S*) stereoisomer.

### QM/MM simulations reveal preferential deacylation of the meropenem Δ2 tautomer

Crystallographic resolution of the KPC-2 carbapenem acyl-enzymes in the Δ1 form was unexpected, given previous reports that such complexes of other class A SBLs are hydrolytically inert^10, 42-44^ and our observation of the meropenem acyl-enzyme of the related carbapenemase SFC-1 (trapped using a Glu166Ala mutation) in the Δ2 form.^45^ Thus, to investigate the behavior of the carbapenem-derived KPC-2 acyl-enzyme in both the Δ1 and Δ2 forms, the KPC-2^E166Q^:meropenem acyl-enzyme structure was used as a starting point for molecular dynamics simulations. In order to simulate native KPC-2, Gln166 was replaced with Glu *in silico*, a change that showed no effect upon the dynamics of the uncomplexed enzyme in triplicate 100 ns MM MD simulations (**Figures S7 and S8)**.^11^ The Δ2-pyrroline (enamine) meropenem-derived acyl-enzyme was modelled using the X-ray structure of the Δ1-(2*R*)-pyrroline (imine) as a guide. In total, four systems (KPC-2^E166Q^:meropenem-Δ2, KPC-2:meropenem-Δ2, KPC-2^E166Q^:meropenem-Δ1-(2*R*) and KPC-2:meropenem-Δ1-(2*R*)) were evaluated in 500 ns triplicate MM MD simulations (for a total of 1.5 μs of MD for each system). No substantial differences between the different acyl-enzymes in overall root mean square deviation (RMSD) or root mean-square fluctuation (RMSF) values were observed for the duration of the simulations; and interactions of the acyl-enzyme C-7 carbonyl with the oxyanion hole and between Glu166 and Asn170 were similarly maintained as described in **Figures S7 - S9** and **Supporting Note 1**. Specifically, during triplicate 500 ns simulations of the KPC-2:meropenem-derived acyl-enzymes, the acyl-enzyme carbonyl remained within the oxyanion hole, with the exception of 1 ns (for each individual repeat) of simulations of the Δ2 tautomer. The observation that the KPC-2:meropenem acyl-enzyme carbonyl is stable within the oxyanion hole of the KPC-2 active site is consistent with the carbapenemase activity of KPC-2 (**Figure S9**) and contrasts with observations of carbapenem acyl-enzymes of other class A enzymes that lack efficient carbapenem-hydrolyzing activity^9, 43, 46^. The 6α-hydroxyethyl group adopts very similar positions in the (energy minimized and equilibrated) starting structures for simulations of the KPC:meropenem acyl-enzymes in both the Δ1 and Δ2 configurations (**Figure S10**); however, in both the MM and QM/MM (see below) simulations the 6α-hydroxyethyl group samples a greater range of dihedral angles in the Δ2 acyl-enzyme, accessing an orientation with a dihedral of c. -70° that is not populated in simulations of the Δ1 tautomer (**Figure S10**).

QM/MM simulations, using as starting structures the KPC-2 meropenem acyl-enzyme structures prepared for the molecular dynamics simulations above, were then employed to investigate the energetics of deacylation of the KPC-2:meropenem acyl-enzymes in both the Δ1-(2*R*) and Δ2-pyrroline forms. (The Δ1-(2*S*) tautomer was not simulated as this form was not observed in either the meropenem or ertapenem acyl-enzyme complexes, and NMR studies of KPC-2 catalyzed carbapenem hydrolysis identified the Δ1-(2*R*) tautomer as formed preferentially.^47^) As in our previous work,^11^ experiments focused on the first stage of deacylation, namely addition of water to the acyl-enzyme carbonyl to form a tetrahedral deacylation intermediate. This first step was simulated as it represents the likely rate-limiting step for the deacylation reaction, and previous simulations accurately differentiated between class A carbapenemases and class A enzymes that are inhibited by carbapenems. ^17^

Tetrahedral intermediate (TI) structures were obtained from conventional umbrella sampling simulations of the 2D MFEP for meropenem deacylation (QM region defined in **Figure S2**) with reaction coordinates defined as described above and in **Supporting Note 1**. The recently introduced adaptive string method (ASM)^20, 48^ finds the minimum free energy pathway (MFEP), “on-the-fly”, for reactions in the space of a set of collective variables, the MFEP is then used to define a path collective variable that serves as a RC in Umbrella Sampling to obtain a free energy profile of the process. Both conventional umbrella sampling and the ASM were implemented to determine the MFEP, and associated Gibbs free energy barriers, for the first stage of deacylation of the meropenem acyl-enzymes in the Δ2 and Δ1-(2*R*) configurations. These ASM calculations identified a clear difference between the two tautomers with respect to the energetics of tetrahedral intermediate formation, with the Δ2 tautomer undergoing this reaction much more readily (ΔG^‡^= 12.30 ± 3.47 kcal/mol) than the Δ1-(2*R*) tautomer (ΔG^‡^= 19.44 ± 1.57 kcal/mol) **Figure S11, Table S3**. Umbrella sampling gave similar results (**Table S3**), but ASM better differentiated between the two carbapenem tautomers and their impacts on the MFEP. The lower, i.e. closer to values derived from experimental measurements of carbapenem turnover, Gibbs Free Energy barrier of reaction calculated by the ASM, compared to conventional umbrella sampling, indicates that the ASM method is better able to optimize the MFEP and provides better description of both the transition state (TS) and overall reaction compared to conventional umbrella sampling.

### Arrangement and stability of H-bond networks in the deacylation transition state

To investigate the basis for the clear differences in the energetics of tetrahedral intermediate formation between the Δ1-(2*R*) and Δ2 pyrroline forms of the KPC-2:meropenem acyl-enzyme, the respective transition states (TS) identified in QM/MM ASM simulations were compared (**Figures 6, S12 and Table S4**). This revealed differences between the Δ1-(2*R*) and Δ2 pyrroline forms in the H-bonding networks each makes to the KPC-2 active site (**Figures 6, S12 and Table S4**). For this analysis, H-bonds were defined by distance and angle, calculated by CPPTRAJ between the protein and meropenem in MFEP calculations.^30^

**Figure 6.**
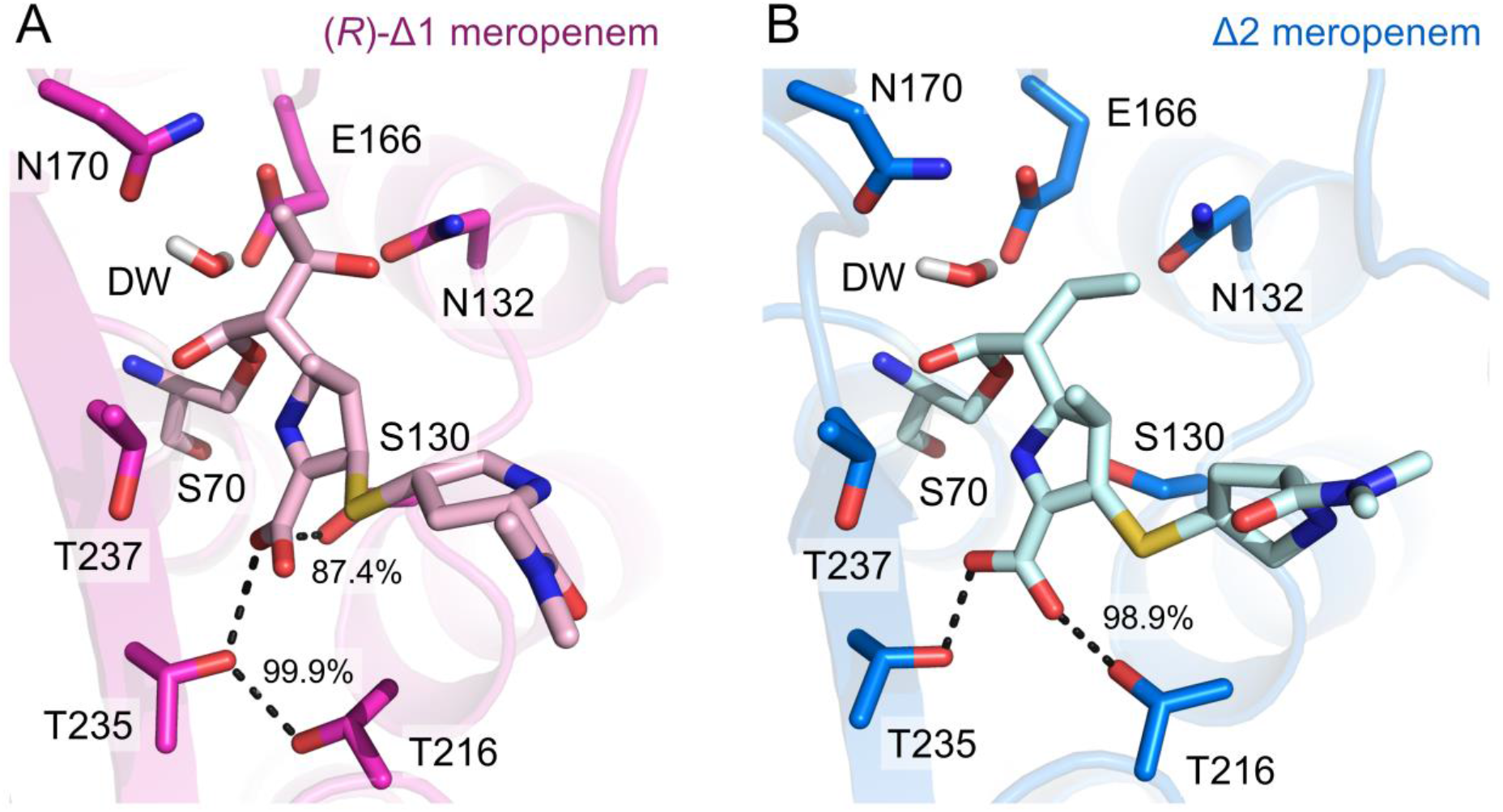
ASM calculations of meropenem tetrahedral intermediate formation reveal changes in hydrogen bonding networks around the C-3 carboxylate of KPC-2:meropenem transition states. Snapshots taken of **A**. the KPC-2:Δ1-meropenem and the **B**. KPC-2:Δ2-meropenem showing the proximity of the deacylating water (DW) proton to E166 (proton transfer) and the hydroxyl to C-7 carbonyl (nucleophilic attack) of meropenem, the transition state (TS) before formation of the tetrahedral intermediate (TI). Key differences in the hydrogen bond network between the protein main chain and the meropenem C-3 carboxylate are shown as dashes. These are labelled with the corresponding % of H-bonds formed over the trajectory.

Both tautomers share common H-bonds to KPC-2, specifically between the acyl-enzyme carbonyl (MER O27) and the backbone amides of Ser70 and Thr237 that form the oxyanion hole, and between Ser70 OG and the side chain Nζ of Lys73. Additional H-bonds are however tautomer-specific (**Figures 6, S12 and Table S4**); the meropenem C-3 carboxylate in the Δ2 tautomer predominantly makes H-bonds to the side chains of Thr216 (98.89 % of frames) and Thr235 (99.58 % of frames), but to those of Thr235 (99.9% of frames) and Ser130 (87.36% of frames) in the Δ1-(2*R*)-derived structure.

Over the entire MFEP ASM calculation, the Δ1-(2*R*) and Δ2 tautomers also differ with respect to H-bonding patterns involving the DW. In simulations of deacylation of the KPC-2:Δ1-(2*R*)-meropenem acyl-enzyme, H-bonds to the DW were promiscuous and involved either the Oδ1 (64.86%) or Nδ2 (6.12%) atoms of Asn170 and the Oε1 (22.63%) or Oε2 (51.54%) atoms of Glu166 (**Table S6**). By comparison, the KPC-2:Δ2-meropenem complex had more consistent H-bond donors/acceptors to the DW in the simulations; H-bonds to Asn170 occurred most often with Oδ1 (74.8%, only 0.26% with Nδ2) and to Glu166 with Oε1 (75.41%, 5.81% with Oε2) (**Table S6**). H-bonds to the DW were not only more consistent in the KPC-2:Δ2-meropenem QM/MM simulations, but were also 10% more frequent in the MFEP calculations (**Table S6**). Hence, in simulations with meropenem in the Δ1-(2*R*) configuration, residues Glu166 and Asn170, as well as the DW, sampled more conformational space, and participated in a reduced number of interactions, compared to the Δ2 tautomer. These findings align with calculations of the DW to C-7 angle (i.e. the angle of nucleophilic attack for deacylation), in which the Δ2 tautomer sampled the favored Bürgi-Dunitz angle^49^ (i.e. between 107° and 109°) in a further 3.42% of simulation frames compared to Δ1 (**Figure S12**). In the Δ2-enamine acylenzyme complex, the DW therefore adopts a more favored orientation for proton transfer to Glu166, and subsequent reaction with the C-7 carbonyl carbon. In contrast, in the Δ1-(2*R*)-imine configuration the DW is less well stabilized, spending larger amounts of the simulation time in positions that are disfavored for either nucleophilic attack or proton transfer events.

## Discussion

KPC-2 efficiently hydrolyzes a wide range of β-lactam classes, including carbapenems with varying C-2 substituents (**Figure 2**), and is of increasing global importance as a cause of antibiotic resistance. Our prior studies of penem (faropenem)^39^ and cephalosporin^11^ (ceftazidime and cefotaxime) turnover by KPC-2 revealed that capture and accommodation of diverse substrates is facilitated by the mobility of loops around the active site. However, the stability of the Ω-loop, and by consequence correct positioning of Glu166, is critical for efficient catalysis. In particular, ceftazidime can bind through disruption of the Ω-loop conformation, with MD simulations indicating the possibility of large movements in this region in the ceftazidime acyl-enzyme, and the potential for alternative conformations of Glu166 that positioned its side chain either ‘in’ to or ‘out’ of the active site. A dual in/out conformation of residue 166 was also observed in a crystal structure of the faropenem-derived acyl-enzyme. In the ‘in’ conformation, Glu166 would be correctly positioned to activate the water molecule for deacylation, whereas the ‘out’ conformation does not promote hydrolysis. These findings explain the relatively poor turnover rates (*k*_cat_) shown by KPC-2 towards ceftazidime and faropenem (**Table 1**). In the carbapenem complexes presented here, disruption of the Ω-loop is less apparent, with in each case the loop being modeled in a single conformation in which all residues are resolved. However, analysis of the B-factors of the Ω-loop (residues 165 to 170) in all KPC-2^E166Q^ acyl-enzyme complex structures obtained to date (with ampicillin, cefalothin, cefotaxime, imipenem, meropenem, ertapenem and faropenem) revealed a negative correlation with steady-state *k*_cat_ values (**Figure 7, Table 1**), indicating that faster antibiotic turnover rates are facilitated by a lower average mobility of the Ω-loop. In particular, the stability and rigidity of the conformations of Glu166 and Asn170 are crucial for productive interactions with the deacylating water molecule, and for keeping conserved, catalytically important active site H-bonding networks intact. Previously determined fast rate constants for carbapenem deacylation^36^ can therefore be attributed to the stability of the Ω-loop in the acylenzyme complexes.

**Figure 7.**
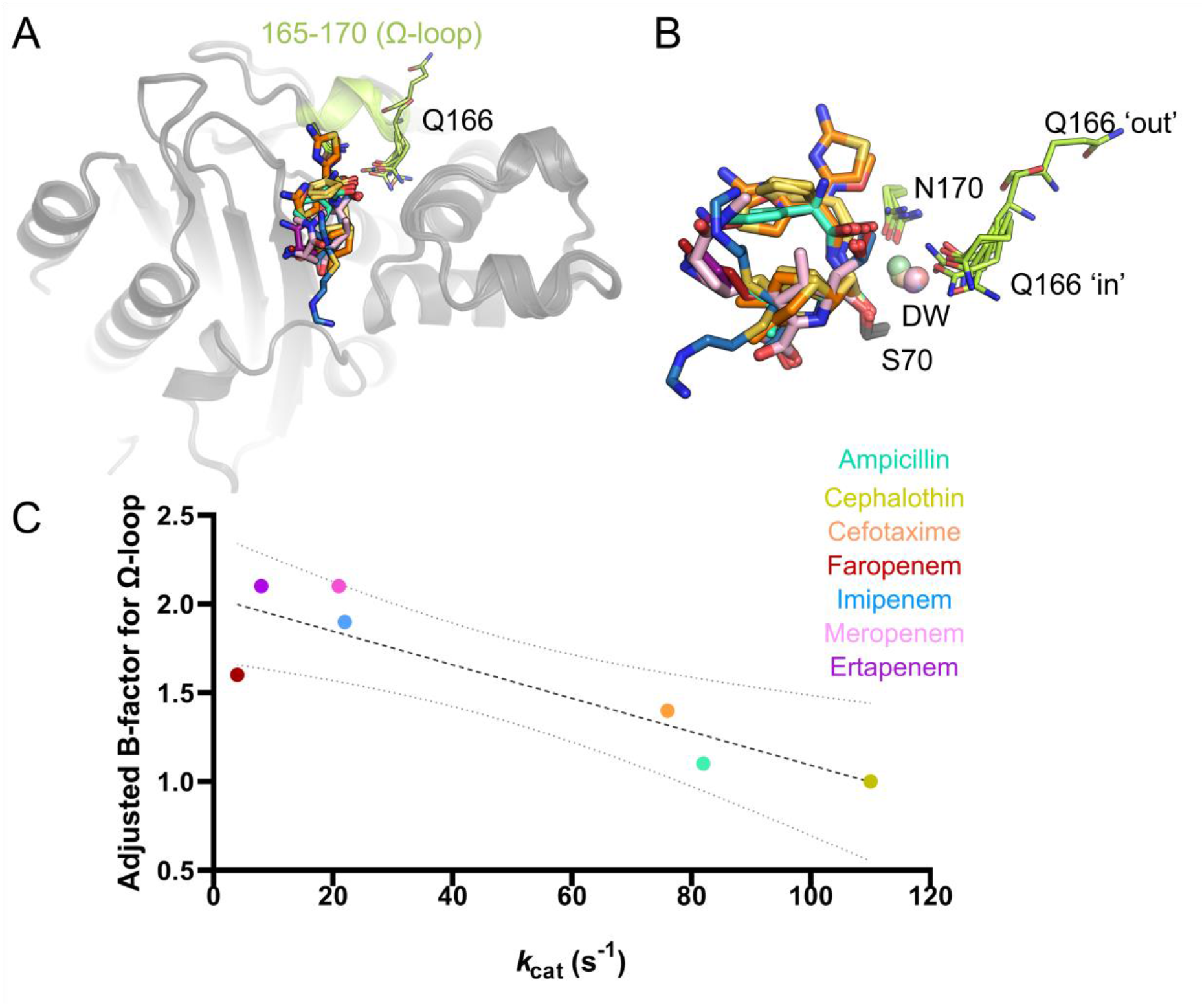
B-factors of the KPC-2 Ω-loop for β-lactam acyl-enzyme complex structures correlate with *k*_cat_. **A** overall view of KPC-2 active site with antibiotic acyl-enzymes, residues 165-170 of the omega loop (highlighted in lime). **B**. Zoom in on active site positions 170 and 166, the position of the water in the putative deacylating position (DW) and corresponding antibiotic acyl-enzymes. Antibiotic-derived acyl-enzymes and DW colored as panel **A. C**. Values for adjusted Ω-loop B-factor and *k*_cat_ from **Table S5** plotted and fit to linear trendline with R^2^ of 0.77. Dotted lines represent 95% confidence intervals for the linear regression fit. Adjusted B-factors were calculated as a ratio vs the average protein B-factor within the crystal structure, see **Table S5**.

X-ray crystal structures are now available for carbapenem acyl-enzymes of multiple class A SBLs (**Figure S13**). These enable comparison of carbapenem binding across a panel of class A β-lactamases which include carbapenemases (SFC-1^18^, GES-5^43^ and KPC-2^50^) and carbapenem-inhibited enzymes (TEM-1^42^, SHV-1^46^, GES-1^43^ and BlaC^51^). Several features of carbapenem binding have been deemed to be important to efficient turnover,^5^ including positioning of the C-7 carbonyl group within the oxyanion hole. In carbapenem acyl-enzyme complexes of both TEM-1 and SHV-1 there is evidence that the acyl-enzyme carbonyl group can ‘flip’ out of this site, destabilizing both this and related structures (e.g. the deacylation tetrahedral intermediate) and in consequence retarding deacylation.^42, 46^ In contrast, in the structures that we report here the acyl-enzyme carbonyl is invariably positioned in the oxyanion hole, and only very transiently moves outside it in the extensive associated molecular simulations.

Efficient carbapenem deacylation has also been associated with orientation of the 6α-hydroxyethyl substituent to prevent interactions with the deacylating water molecule DW that have been proposed to reduce its nucleophilicity.^18^ The structures reported here support this conclusion, with this group consistently positioned to interact with Asn132 and lying beyond H-bonding distance of DW. Both MM MD and ASM simulations nevertheless identify that conformations closer to DW are occasionally sampled by the meropenem acyl-enzyme (**Figure S10**), consistent with recent reports demonstrating that the imipenem acyl-enzyme complex of a KPC-2 F72Y mutant can form an H-bond from the 6α-hydroxyethyl to DW. However, the low frequency (occurring in only 4.71 % and 1.31 % of frames in the Δ2 and Δ1-(2*R*) ASM simulations, respectively) with which these contacts occur in the simulations support the conclusion that reduced interaction of the 6α-hydroxyethyl group with DW remains an important contributor to carbapenem-hydrolyzing ability on Class A enzymes.

Recently, use of the N170A mutant has enabled hydrolyzed antibiotics to be non-covalently trapped in the KPC-2 active site, yielding structures of KPC-2^N170A^:ampicillin (PDB:6XD7, 1.65 Å resolution) and KPC-2^N170A^:imipenem (Δ2) (PDB:6XJ8, 2.05 Å resolution)).^36^ When considered alongside the KPC-2^E166Q^ acyl-enzyme complexes reported here, the combined data may represent how the enzyme accommodates substrates during and after deacylation events (**Figure S14**). Deacylation of the ampicillin-derived acyl-enzyme results in changes to the active site hydrogen bonding network, with the newly-formed C-7, and C-3, carboxylates of the non-covalently bound product interacting respectively with the Thr237 oxygen/Ser70 backbone amides and Ser70 OG oxygen (**Figure S14**). The C-6 substituent also moves out into solvent, indicating a system that is primed for product release. On the other hand, the non-covalent imipenem-derived product retains the same general positioning within the KPC-2 active site, including interactions involving the C-3 carboxylate, that are observed in the imipenem acyl-enzyme structure presented here (**Figure S14**) and in the equivalent complex with the F72Y mutant. These differences may reflect differing affinities of KPC-2 for ampicillin and imipenem, and by implication their respective hydrolysis products, that are reflected in their differing *K*_M_ values (**Table 1**).

The efficiency of carbapenem hydrolysis by SBLs is also thought to be affected by the ability of the carbapenem-derived products to tautomerize between the Δ1-(2*R*/2*S*) and Δ2 forms (**Figure 3**). Our previous NMR solution studies identified formation of the Δ2 tautomer by multiple carbapenemases, followed by rapid tautomerization to the Δ1-(2*R*) form.^47^ Similarly, kinetic^52-53^ and Raman^10^ studies on carbapenem-inhibited enzymes (i.e. TEM-1 and SHV-1) indicated initial formation of the Δ2 tautomer, which subsequently either deacylates or isomerizes to the Δ1 form(s). The present data, which crystallographically identify acylenzymes of KPC-2 with multiple carbapenems as all in the Δ1-(2*R*) form, provides evidence that this isomerization can also take place in carbapenem-hydrolyzing enzymes. Specifically, our high-resolution crystal structure of the KPC-2:imipenem-derived acyl-enzyme reveals that both (2*R*) and (2*S*) stereoisomers of the Δ1 tautomer can exist at the same time, suggesting that, at least in this system, tautomer interconversion has a low energy barrier and can occur on (crystallized) KPC-2 after initial formation of the Δ2 acyl-enzyme (**Figure 3**). In previous investigations of carbapenem-inhibited enzymes, accumulation of the Δ1 acyl-enzyme as a stable species led to its designation as (near) hydrolytically inert. Consistent with these observations, our QM/MM calculations (using the adaptive string method) modeling formation of the tetrahedral deacylation intermediate show that the Δ1-(2*R*) tautomer of meropenem presents a significantly larger barrier for acyl-enzyme deacylation by KPC-2 than does the Δ2 form (19.44 ± 1.57 kcal mol^-1^ compared to 12.30 ± 3.47 kcal mol^-1^, **Table S3**).

The likely basis for differences in the susceptibility to hydrolysis of the Δ1 and Δ2 tautomers is explained by analysis of the QM/MM simulations with the adaptive string method. Previous studies indicate that the 6α-hydroxyethyl substituents of carbapenem acyl-enzymes can adopt multiple positions and orientations across different class A β-lactamases.^5, 18, 42, 46^ This enables interactions with either Glu166 (as seen in the BlaC:doripenem acyl-enzyme, PDB 3IQA, **Figure S13**^42^) or the deacylating water (SHV-1:meropenem acyl-enzyme, PDB 2ZD8, **Figure S13**^46^). Our previous MD study of the efficient carbapenemase SFC-1 established that the meropenem 6α-hydroxyethyl group established consistent H-bond interactions with Asn132,^18^ preventing contacts with the deacylating water that would reduce its nucleophilicity and slow deacylation, and explaining the basis for SFC-1 carbapenemase activity.^18^ In this work, QM/MM simulations revealed further contributors to carbapenem hydrolysis, as evident in comparisons of the transition states of the meropenem-derived tautomers.

Specifically, hydrogen bonding around the C-3 carboxylate (**Figure 6**) requires mutually exclusive networks in the Δ1-(2*R*) and Δ2 tautomer complexes, that involve residues Thr216, Thr235 and Ser130, and result in the C-3 carboxylate adopting different orientations in the two meropenem tautomers. In consequence, the N-4 N-H group of the Δ2 meropenem (enamine) tautomer will be electrostatically stabilized by proximity to the O-9 oxygen. Furthermore, the proximity of the enamine N-4 to the acyl-enzyme ether OG and carbonyl provides for further electrostatic interaction, which will strengthen during formation of the deacylation transition state and tetrahedral intermediate (**5, Figure 1**) as the nascent oxyanion accumulates negative charge. The effect will be selective stabilization of these species, relative to the starting acylenzyme, with consequent reduction in the free-energy barrier for deacylation. In contrast, the unprotonated imine N-4 of the Δ1-(2*R*) tautomer will experience no such stabilizing effect, indeed the proximity of its lone pair to Ser70 OG is expected to be relatively destabilizing for tetrahedral intermediate formation.

In addition, the two tautomers differ in the respective interaction networks that involve the deacylating water molecule (DW). In simulations of the Δ2 tautomer the DW makes consistent H-bonds to specific atoms in the Asn170 and Glu166 side chains, whilst for the Δ1-(2*R*) tautomer interactions of DW with its H-bonding partners are more variable. Therefore, in the Δ2-enamine form of the KPC-2:meropenem acyl-enzyme, the position and orientation of the DW in the transition state is more constrained and more stable, facilitating ready addition to the acyl-enzyme carbonyl. Taken together with the stabilizing contributions of charge-based interactions involving protonated N-4 in the deacylation transition state and intermediate, this is expected to result in a lower free energy barrier for tetrahedral intermediate formation, hence promoting deacylation.

Our crystallographic observation of the KPC-2:carbapenem acyl-enzymes in the Δ1 configuration might be considered surprising in light of this being regarded as a hydrolytically inert species, and the expectation that its formation might be disfavored or prevented in carbapenem-hydrolyzing Class A enzymes such as KPC-2. Given the evidence that Δ1-tautomers are more stable than the Δ2-form^47, 54^ it may be that the former accumulate *in crystallo*, where deacylation is slower than in solution, perhaps due to intermolecular interactions between molecules in the crystalline lattice. Another possibility is that Class A carbapenemases might accelerate deacylation of the Δ1 acyl-enzyme, such that it no longer accumulates as an inhibitory species. Our data suggest however that this is not the case, that is the carbapenem-hydrolyzing activity of KPC-2, and by implication other Class A enzymes with carbapenemase activity, arises primarily from a favorable rate of deacylation of the Δ2 acylenzyme, as evidenced by the low barriers obtained from QM/MM calculations, rather than an ability to prevent formation of the Δ1 tautomer or accelerate its deacylation. A further question is then how KPC-2 and related enzymes avoid the accumulation of the Δ1 acyl-enzyme in solution, that our QM/MM calculations identify as resistant to deacylation. In this respect our crystallographic observation of the imipenem acyl-enzyme in both Δ1-(2*R*) and Δ1-(2*S*) configurations is significant, as it gives rise to the possibility that these species are in exchange in the enzyme-bound form, i.e. that tautomerization, *via* the Δ2 form, is freely reversible. Thus, efficient carbapenem turnover is possible as the acyl-enzyme retains access to a state (Δ2) for which it is deacylation-competent, even though the Δ1 forms remain hydrolytically inert.

## Conclusions

Taken together, the extensive new structural data and molecular simulations presented here provide the most detailed picture to date of β-lactam antibiotic hydrolysis by KPC-2. Our data show that changes in active site architecture, particularly of the Ω-loop, are crucial determinants of the rate of antibiotic hydrolysis by this versatile enzyme. Although mobility of the Ω-loop may be required for the broad substrate selectivity of KPC-2, it can negatively impact turnover, with the stability and correct positioning of Glu166 essential for efficient hydrolysis. ASM calculations, based upon the crystal structures reported here, further revealed that the meropenem acyl-enzyme in the Δ2 tautomer is efficiently deacylated compared to the Δ1-(2*R*) form, a proposal consistent with solution analyses of products of β-lactam hydrolysis by both SBLs and MBLs. Exclusive electrostatic interactions that differ between tautomers, particularly those involving the C-3 carboxylate, and the less stable interactions made by the deacylating water (DW) in the meropenem Δ1-(2*R*) acyl-enzyme, compared to the Δ2 tautomer, contribute to a 7.1 kcal/mol increase in activation free energy barrier for deacylation. ASM simulations therefore have the sensitivity to identify the differing sets of interactions in the deacylation-competent (Δ2) and deacylation-incompetent (Δ1-(2*R*)) forms of the KPC-2:meropenem acyl-enzyme, that respectively promote or prevent carbapenem turnover, and so discriminate between closely related complexes that nevertheless differ in reactivity. The results thus demonstrate the utility of the ASM method as a tool with which to investigate the energetics of enzyme-catalyzed reactions, and suggest that modifying β-lactams to promote formation of Δ1-type acyl-enzyme complexes represents one possible route to overcoming β-lactam resistance caused by Class A carbapenemases.

## Acknowledgements

Research was supported by the Biotechnology and Biological Sciences Research Council (SWBioDTP [BB/J014400/1], studentship to C. L. T. and BB/T008741/1, studentship to M.B.). C. L. T, J. S, A. J. M and C. J. S thank the Medical Research Council (MR/T016035/1). C. J. S. thanks the Medical Research Council and the Wellcome Trust for funding. A.J.M. thanks the U.K. Engineering and Physical Science Research Council (EPSRC grant no. EP/M022609/1) for support. This work is part of a project that has received funding from the European Research Council under the European Horizon 2020 research and innovation programme (PREDACTED Advanced Grant Agreement no. 101021207) to A.J.M. X-ray diffraction data were collected at the BL13–XALOC beamline at the ALBA Synchrotron with the collaboration of ALBA staff. We also thank Diamond Light Source for beamtime (proposals 172122 and 23269) and the staff of beamlines I24 and I04 for assistance. This work was carried out using the computational facilities of the *Advanced Computing Research Centre*, University of Bristol - http://www.bristol.ac.uk/acrc/.

## Data availability

For all crystal structures presented herein, atomic coordinates and structure factors have been deposited to the Worldwide Protein Data Bank (PDB, wwpdb.org) under accession codes 8AKI (KPC-2^E166Q^:ampicillin), 8AKJ (KPC-2^E166Q^:cephalothin), 8AKK (KPC-2^E166Q^:imipenem), 8AKL (KPC-2^E166Q^:meropenem) and 8AKM (KPC-2^E166Q^:ertapenem). Example structures of the acyl-enzyme complex, transition state and tetrahedral intermediate of both tautomers and input files for all simulations are available at the University of Bristol Research Data Repository (https://data.bris.ac.uk/data/). Ligand parameter files are available at XXXX (figshare).

