## Supporting Information for "Tautomer-specific deacylation and Ω-loop flexibility explain carbapenem-hydrolyzing, broad-spectrum activity of the KPC-2 β-lactamase"

**Contents**

**Tables**

Table S1. Data collection and refinement statistics for KPC-2$^{E166Q}$ crystal structures.

Table S2. Ligand statistics from KPC-2 crystal structures presented herein

Table S3. Calculated activation free energy barriers for meropenem derived complex deacylation by KPC-2.

Table S4. H-bonds between the meropenem (MER) acyl-enzyme complex and the KPC-2 protein mainchain observed during transition state frames in QM/MM calculations.

Table S5. Crystallographic B-factor comparisons with $k_{cat}$ values

Table S6. Hydrogen bond interactions observed for of the deacylating water over adaptive string method (ASM) simulations

Table S7. Active site distances in molecular dynamics (MD), umbrella sampling (US) and adaptive string method (ASM) calculations

Table S8 Solvent accessible surface area (SASA) in Å$^2$ for the KPC-2 active site during KPC-2:meropenem simulations

**Figures**

Figure S1. View of the KPC-2 apo-enzyme active site.

Figure S2. QM region (black) used for QM/MM calculations

Figure S3. Evidence for two conformations of the thiophenylacetamido C-7 substituent in the KPC-2$^{E166Q}$:cefalothin complex.

Figure S4. Carbapenem-derived acyl-enzyme complexes colored with respect to atomic B-factor

Figure S5. Possible hydrogen bonding interactions in KPC-2$^{E166Q}$ carbapenem acyl-enzyme complexes

Figure S6. Possible hydrogen bonding interactions in KPC-2$^{E166Q}$ carbapenem acyl-enzyme complexes.

Figure S7. RMSD plots for the KPC-2:meropenem complex over triplicate 500ns simulations

Figure S8. Per residue RMSF plots for the KPC-2:meropenem-derived acyl-enzyme complex over triplicate 500 ns simulations.

Figure S9. Distance analyses over KPC-2:meropenem-derived acyl-enzyme triplicate simulations.

Figure S10. Position of the hydroxyethyl in MM and QM simulations.

Figure S11. ASM calculations for tetrahedral intermediate formation from the meropenem-derived acyl-enzyme in KPC-2 catalysis

Figure S12. Transition state analysis of adaptive string method calculations

Figure S13. Comparisons of the interactions of carbapenem acyl-enzymes within Class A SBLs.

Figure S14. Comparison of KPC-2:antibiotic acyl-enzyme and non-covalent hydrolysis product complexes.

Figure S15. Ser130 interactions throughout MD and ASM simulations.

Figure S16. Lys234 networks throughout ASM simulations

**Supporting notes:**

Supporting note1

Supporting note 2

### Tables

**Table S1. Data collection and refinement statistics for KPC-2$^{E166Q}$ crystal structures**

| | KPC-2$^{E166Q}$: ampicillin | KPC-2$^{E166Q}$: cephalothin | KPC-2$^{E166Q}$: imipenem | KPC-2$^{E166Q}$: meropenem | KPC-2$^{E166Q}$: ertapenem |
|---|---|---|---|---|---|
| **PDB ID** | 8AKI | 8AKJ | 8AKK | 8AKL | 8AKM |
| **Data collection** | | | | | |
| **Space group** | $P2_12_12$ | $P2_12_12$ | $P2_12_12$ | $P2_12_12$ | $P2_12_12$ |
| **Molecules/ASU** | 1 | 1 | 1 | 1 | 1 |
| **Cell dimensions** | | | | | |
| **a, b, c (Å)** | 60.51, 79.38, 56.21 | 60.51, 79.58, 55.95 | 60.66, 79.48, 55.94 | 60.56, 79.21, 56.26 | 60.25, 78.55, 55.59 |
| **α, β, γ (°)** | 90, 90, 90 | 90, 90, 90 | 90, 90, 90 | 90, 90, 90 | 90, 90, 90 |
| **Resolution (Å)** | 48.12-1.40 (1.42-1.40) | 48.17-1.35 (1.37-1.35) | 48.22-1.36 (1.38-1.36) | 45.86- 1.35 (1.37-1.35) | 47.80-1.25 (1.27-1.25) |
| **$R_{pim}$** | 0.034 (0.665) | 0.032 (0.553) | 0.038 (0.987) | 0.030 (0.552) | 0.031 (0.656) |
| **$CC_{1/2}$** | 0.999 (0.613) | 0.999 (0.726) | 0.999 (0.519) | 0.999 (0.858) | 0.999 (0.574) |
| **$I/\sigma(I)$** | 13.5 (1.2) | 14.4 (1.2) | 10.5 (0.9) | 14.7 (1.3) | 12.0 (1.1) |
| **Completeness (%)** | 100 (100) | 99 (97.8) | 100 (100) | 100 (100) | 100 (100) |
| **Redundancy** | 13.1 (13.1) | 13.3 (13.6) | 13.1 (13.2) | 13.1 (13.3) | 12.8 (12.9) |
| **Refinement** | | | | | |
| **Resolution (Å)** | 48.12-1.40 | 48.17-1.35 | 48.22-1.36 | 45.86-1.35 | 47.804-1.25 |
| **No. reflections** | 54,007 | 59,287 | 58,791 | 60,064 | 73,569 |
| **$R_{work}$ / $R_{free}$** | 0.157/0.1849 | 0.1504/0.1776 | 0.1542/0.1797 | 0.1365/0.1662 | 0.1511/0.1642 |
| **No. non-H atoms** | | | | | |
| **Protein** | 2094 | 2126 | 2125 | 2059 | 2119 |
| **Solvent** | 320 | 281 | 301 | 300 | 293 |
| **Ligand** | 24 | 44 | 40 | 52 | 66 |
| **B-factors** | | | | | |
| **Protein** | 18 | 17 | 19 | 18 | 18 |
| **Solvent** | 39 | 35 | 40 | 37 | 38 |
| **Ligand** | 17 | 19 | 30 | 23 | 34 |
| **R.m.s. deviations** | | | | | |
| **Bond lengths (Å)** | 0.009 | 0.009 | 0.008 | 0.008 | 0.007 |
| **Bond angles (°)** | 1.008 | 1.043 | 0.994 | 0.977 | 1.021 |
| **Ramachandran (%)** | | | | | |
| **Outliers** | 0.00 | 0.00 | 0.00 | 0.00 | 0.00 |
| **Favored** | 98.87 | 98.87 | 98.50 | 98.13 | 98.50 |

**Values in parentheses are for the outer shell.**

**Table S2. Ligand statistics from KPC-2 crystal structures presented herein**

| Ligand | B-Factors | Occupancies | RSCC (PDB) |
|---|---|---|---|
| Ampicillin | 17 | 1.0 | 0.97 |
| Cefalothin | 19 | 0.09/0.91 | 0.96/0.96 |
| Imipenem | 30 | 0.5/0.5 | 0.94/0.94 |
| Meropenem | 23 | 1 | 0.89 |
| Ertapenem | 34 | 1 | 0.96 |

**Table S3. Calculated activation free energy barriers for meropenem derived complex deacylation by KPC-2.**

| | **Adaptive string method** | | **Umbrella sampling** | |
|---|---|---|---|---|
| | **Δ1** | **Δ2** | **Δ1** | **Δ2** |
| $\Delta^{\ddagger}\boldsymbol{G}_{\text{calc}}$ | 19.44 (1.57) | 12.30 (3.47) | 23.08 (1.14) | 19.67 (1.12) |

Standard deviations are shown in parentheses

**Table S4. H-bonds between the meropenem (MER) acyl-enzyme and KPC-2 during transition state frames in QM/MM calculations.** Main differences between tautomers are highlighted in green.

| **Acceptor atom** | **Donor atom** | **Adaptive string method** | | **Umbrella Sampling** | |
|---|---|---|---|---|---|
| | | **Δ1** | **Δ2** | **Δ1** | **Δ2** |
| **MER O9** | **Thr235 OG1** | 73.77 | 99.58 | 98.3 | 100 |
| **MER O10** | **Thr216 OG1** | - | 98.89 | - | 65.8 |
| **MER O27** | **Thr237 N** | 79.04 | 77.54 | 94.2 | 79.2 |
| **MER O27** | **S70 N** | 93.46 | 84.18 | 100 | 98.3 |
| **MER O9** | **Ser130 OG** | 87.36 | - | 83.3 | 5.8 |
| **MER O13** | **Asn132 ND2** | 44.06 | 39.56 | 43.3 | 49.2 |
| **Asn 132 OD1** | **MER O13** | - | 9.87 | - | 3.33 |
| **MERO9** | **Thr237 OG1** | - | 10.83 | - | 33.3 |
| **MER O10** | **Thr235 OG1** | 26.14 | 0.35 | - | - |
| **MER O9** | **Lys234 NZ** | 6.74 | - | - | - |
| **MER N4** | **Ser130 OG** | 6.88 | 0.1 | - | - |

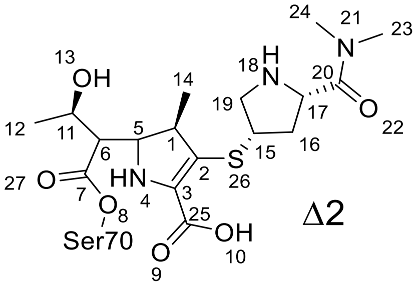


Δ2

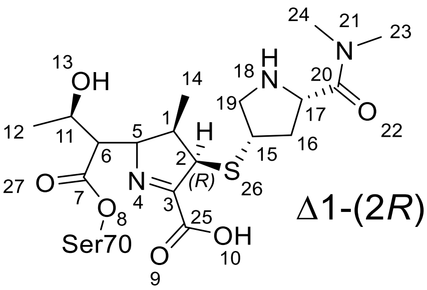


Δ1-(2*R*)

**Table S5. Crystallographic B-factor comparisons with $k_{cat}$ values**

| Antibiotic | $k_{cat}$ ($s^{-1}$) | B-factor (protein) | B-factor (Ω-loop) | Adjusted increase in B-factor (Ω-loop/protein) |
|---|---|---|---|---|
| Ampicillin | 82 | 18 | 21 | 1.1 |
| Cefalothin | 110 | 17 | 17 | 1.0 |
| Cefotaxime | 76 | 24 | 33 | 1.4 |
| Faropenem | 4 | 16 | 26 | 1.6 |
| Imipenem | 22 | 19 | 37 | 1.9 |
| Meropenem | 21 | 18 | 38 | 2.1 |
| Ertapenem | 8 | 18 | 38 | 2.1 |

**Table S6. Hydrogen bond interactions observed for the deacylating water over adaptive string method (ASM) simulations**

| Acceptor Atom | Donor | Trajectory (%) | |
|---|---|---|---|
| | | **Δ1** | **Δ2** |
| Asn170 OD1 | DW | 64.86 | 74.8 |
| Glu166 OE2 | DW | 51.54 | 75.41 |
| Glu166 OE1 | DW | 22.63 | 5.81 |
| DW | Asn 170 ND2 | 6.12 | 0.16 |
| DW | Lys73 NZ | 5.24 | 4.54 |
| | **Total %** | **150.39** | **160.72** |

**Table S7. Active site distances in molecular dynamics (MD), umbrella sampling (US) and adaptive string method (ASM) calculations**

| | | S70-oxyanion carbonyl | T237-oxyanion carbonyl | 166-170 | 132-hydroxyethyl oxygen |
|---|---|---|---|---|---|
| **MD** | **(Δ1)** | 3.05 (0.28) | 2.91 (0.13) | 5.44 (1.07) | 3.28 (0.64) |
| | **(Δ2)** | 2.86 (0.18) | 2.9 (0.24) | 5.63 (0.88) | 3.34 (0.54) |
| **US** | **(Δ1)** | 2.74 (0.07) | 2.84 (0.15) | 4.85 (0.15) | 3.04 (0.23) |
| | **(Δ2)** | 2.79 (0.09) | 2.9 (0.14) | 4.72 (0.19) | 3.32 (0.52) |
| **ASM** | **(Δ1)** | 2.83 (0.24) | 2.85 (0.15) | 4.97 (0.22) | 3.87 (0.46) |
| | **(Δ2)** | 2.92 (0.31) | 2.89 (0.14) | 4.82 (0.26) | 3.7 (0.29) |

Standard deviations are shown in parentheses

**Table S8 Solvent accessible surface area (SASA) in Å$^2$ for the KPC-2 active site during KPC-2:meropenem simulations**

| Simulation | Δ2 | Δ1 |
|---|---|---|
| 1.5 ns MD | 1416.9 (44.3) | 1464.1 (76.4) |
| 900 ps QM/MM | 1454.3 (56.0) | 1408.0 (23.5) |
| US calculations | 1413 (34.4) | 1400 (26.1) |

Standard deviations are shown in parentheses

**Figures**

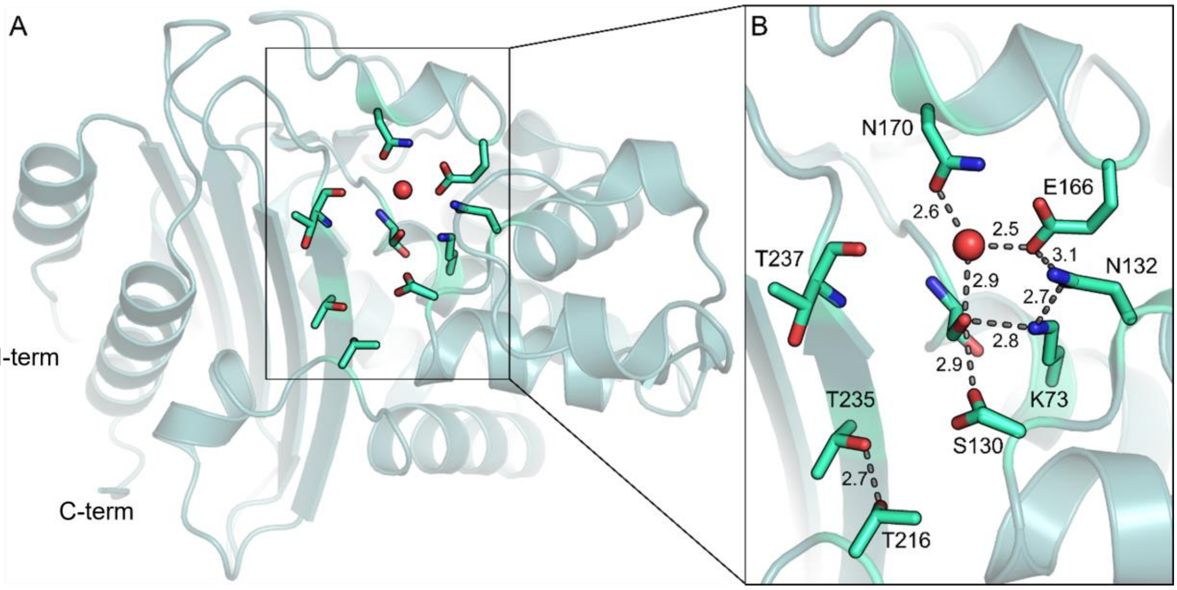


**Figure S1. View of the KPC-2 apo-enzyme active site. A.** Overall structure of KPC-2 (PDB ID 5UL8[1]) shown as teal cartoon with key active site residues as green-cyan sticks. **B.** Zoom in of the KPC-2 active site with key active site residues labelled and apo-enzyme hydrogen bonding networks ≤3.1 Å shown as grey dashes. Networks between residues S70, K73, S130, N132, E166, N170 and the deacylating water (DW, red sphere) are largely conserved across class A serine-β-lactamases.

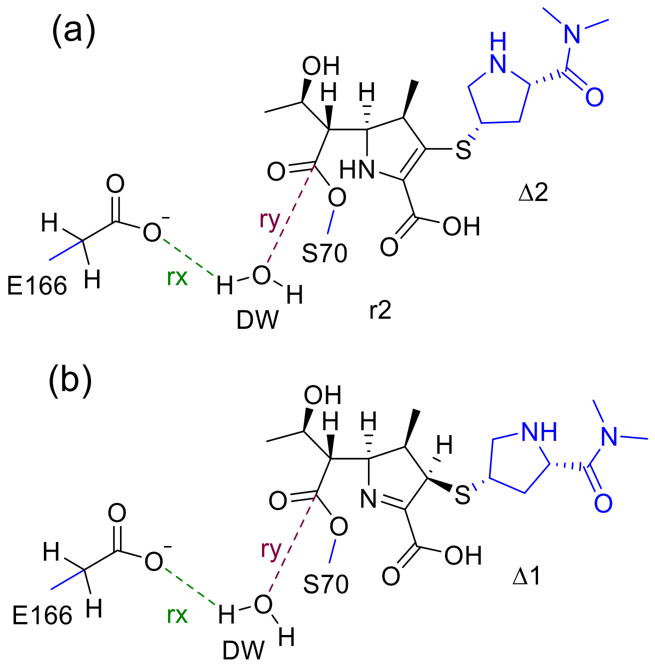


**Figure S2. QM region (black) used for QM/MM calculations.** Acyl-enzyme complexes of **(a)** Δ2 meropenem and **(b)** Δ1-(2*R*) meropenem. The distances between the transferred proton of the deacylating water (DW) and the Glu166 side chain (reaction coordinate 1: rx = d(OεGlu166–HDW) – d(HDW–ODW)) and the distance between the oxygen of the DW and meropenem-derived C-7 carbon (reaction coordinate 2: ry = d(C-7meropenem–ODW)) are labelled in green and red respectively. The collective variables used for the ASM were the distance between the Glu166 side chain and the closest proton of the deacylating water (collective variable 1: cv1 = d(OεGlu166–HDW) (shown in green) and the distance between the oxygen of the DW and meropenem derived C-7 carbon (collective variable 2: cv2 = d(C-7meropenem-ODW) (shown in red).

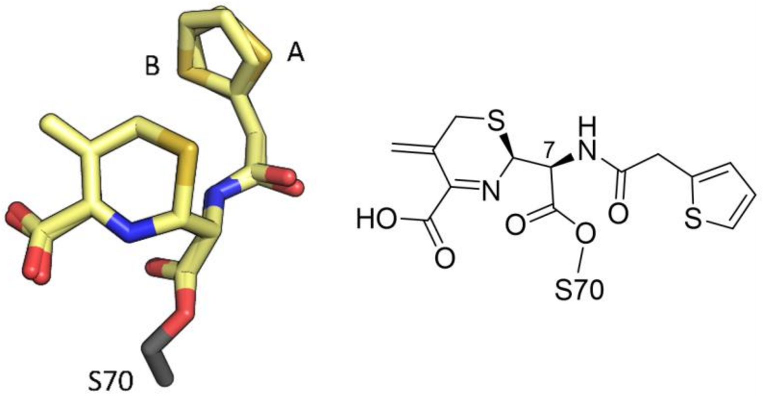


**Figure S3. Evidence for two conformations of the thiophenylacetamido C-7 substituent in the KPC-2$^{E166Q}$:cefalothin complex.** Cefalothin is shown as yellow sticks with the two orientations of the C-7 thiophenylacetamido group labelled as A and B with occupancies of 0.09 and 0.91 respectively, as defined after refinement in Phenix.

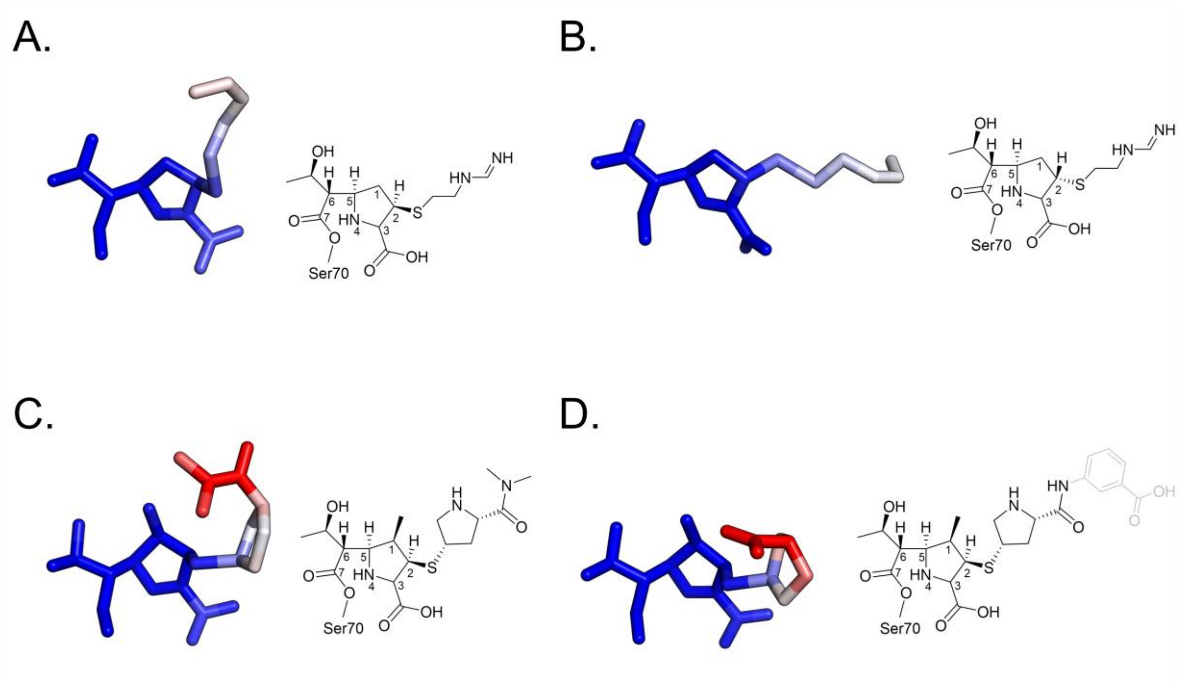


**Figure S4. Carbapenem-derived acyl-enzyme complexes colored with respect to atomic B-factor. A** Imipenem acyl-enzyme complex conformation A (Δ1-2*R*). **B.** Imipenem acyl-enzyme complex conformation B (Δ1-2*S*). **C.** Meropenem acyl-enzyme complex. **D.** Ertapenem acyl-enzyme complex. Atoms highlighted in grey could not be confidently modelled into the experimental electron density, indicating mobility.

**a)**

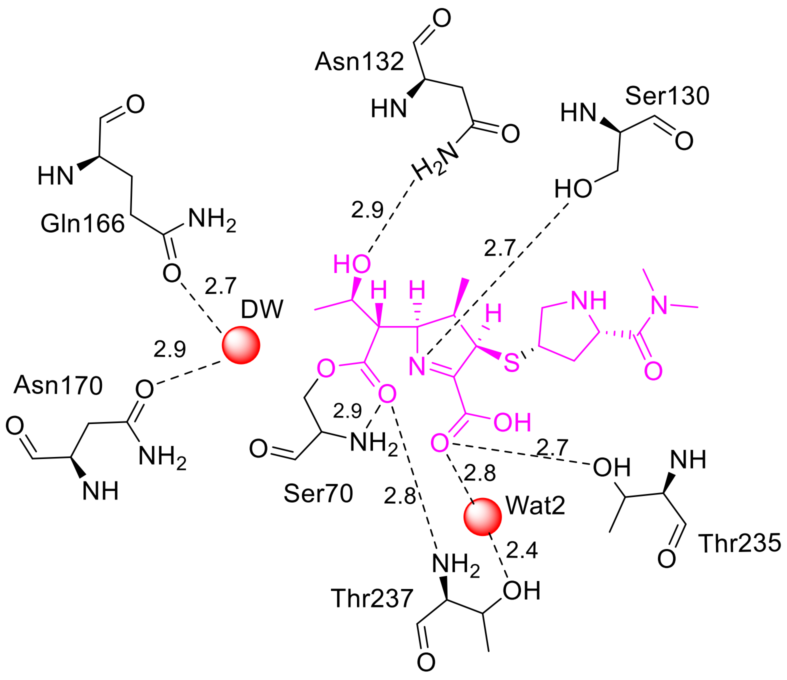


**b)**

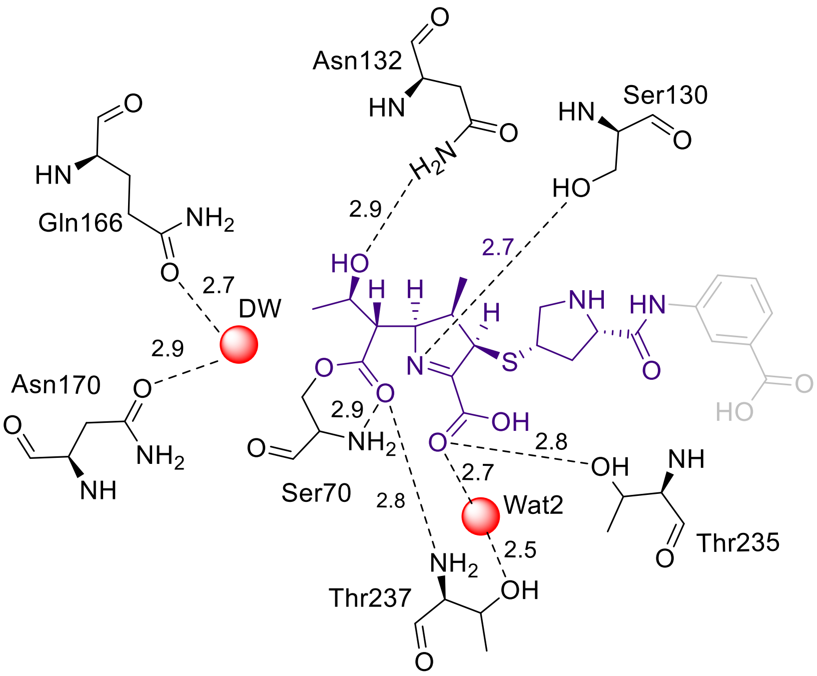


**Figure S5. Possible hydrogen bonding interactions in KPC-2$^{E166Q}$ carbapenem acyl-enzyme complexes. a)** KPC-2$^{E166Q}$:meropenem acyl-enzyme complex, **b)** KPC-2$^{E166Q}$:ertapenem acyl-enzyme complex.

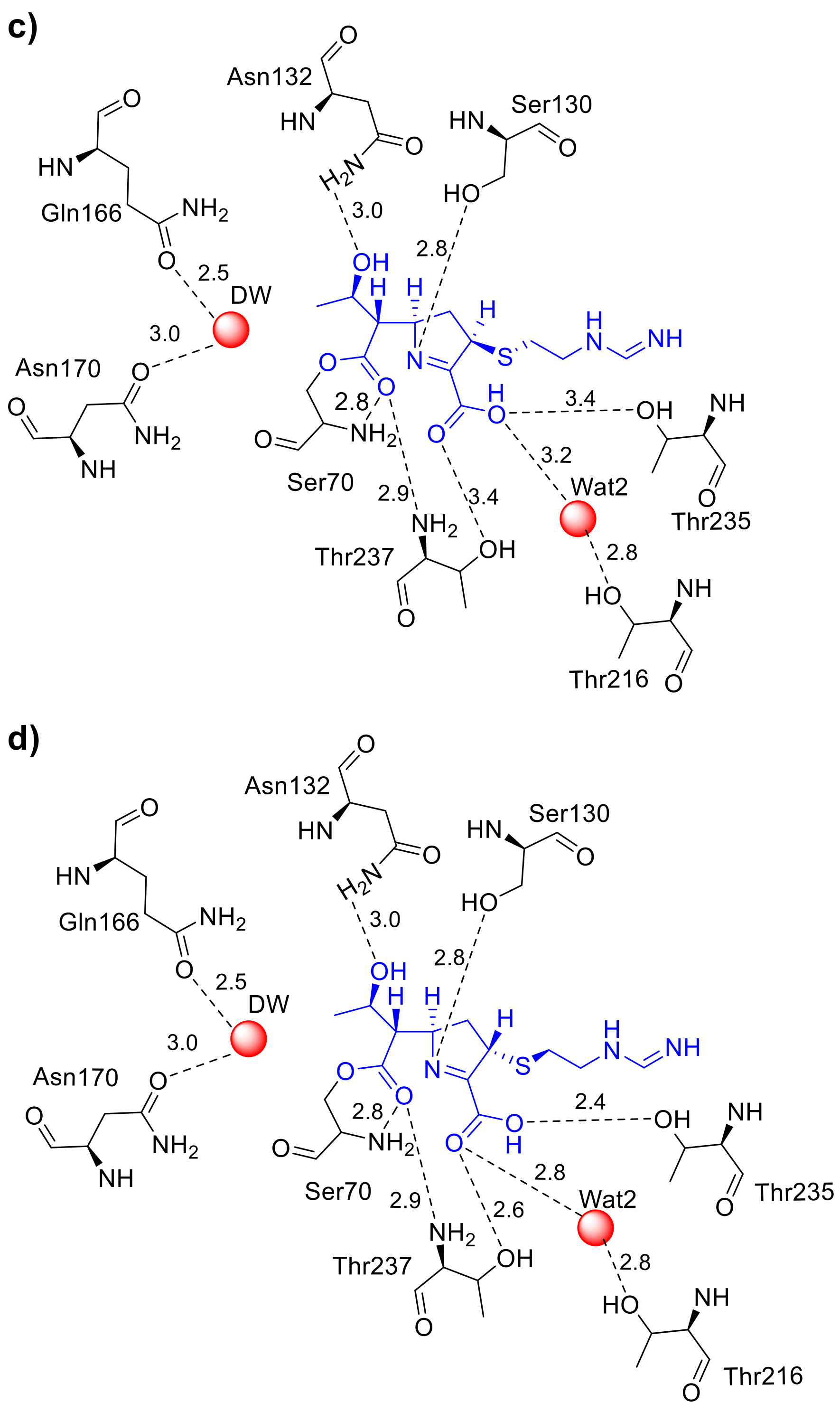


**Figure S6. Possible hydrogen bonding interactions in KPC-2^E166Q^ imipenem acyl-enzyme complexes.** **c)** KPC-2$^{E166Q}$:imipenem (conf A, (2*R*)-configuration) acyl-enzyme complex and **d)** KPC-2$^{E166Q}$:imipenem (conf B, (2*S*)-configuration) acyl-enzyme complex.

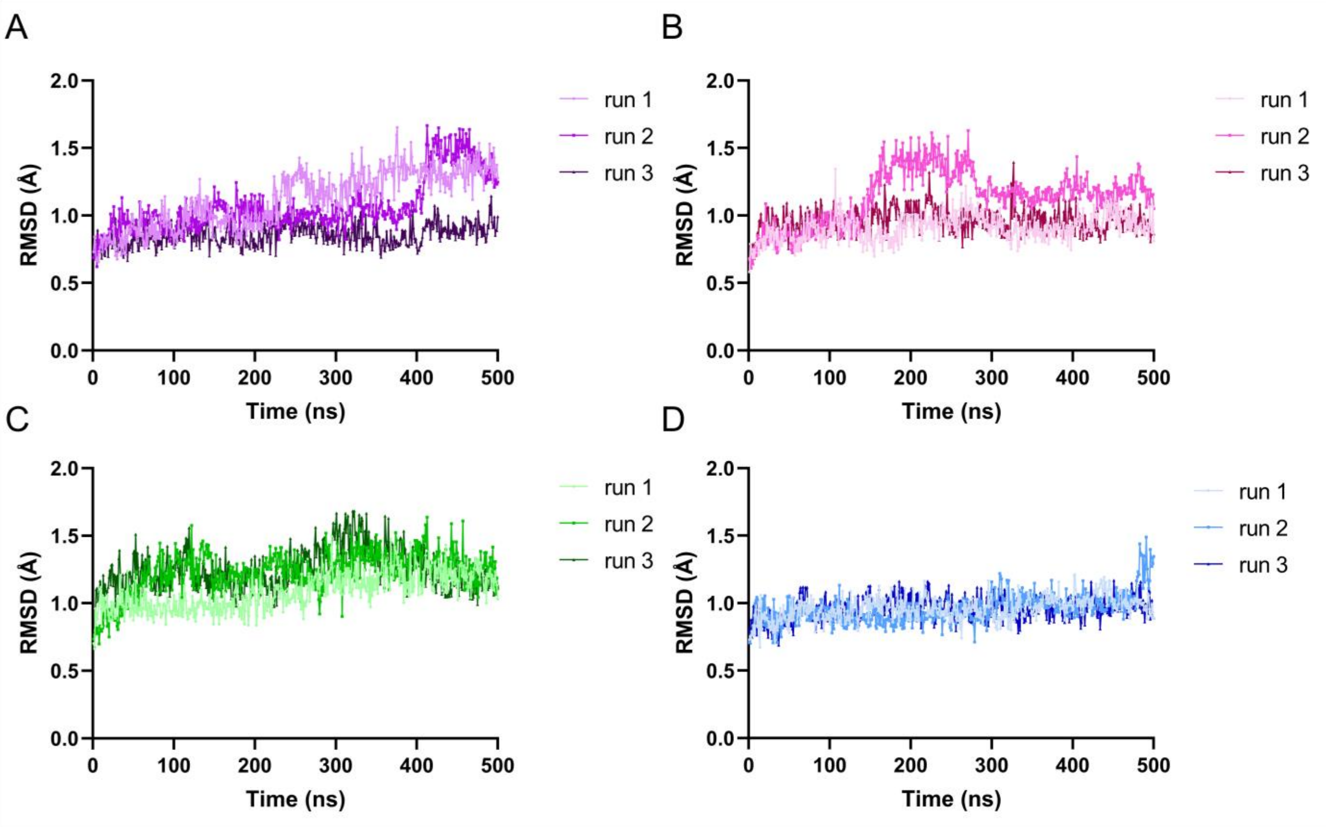


**Figure S7. RMSD plots for the KPC-2:meropenem complex over triplicate 500ns simulations A.** KPC-2$^{E166Q}$:meropenem (Δ1-(2*R*) tautomer), purple. **B.** KPC-2:meropenem (Δ1-(2*R*) tautomer), pink. **C.** KPC-2$^{E166Q}$:meropenem (Δ2 tautomer), green. **D.** KPC-2:meropenem (Δ2 tautomer), blue.

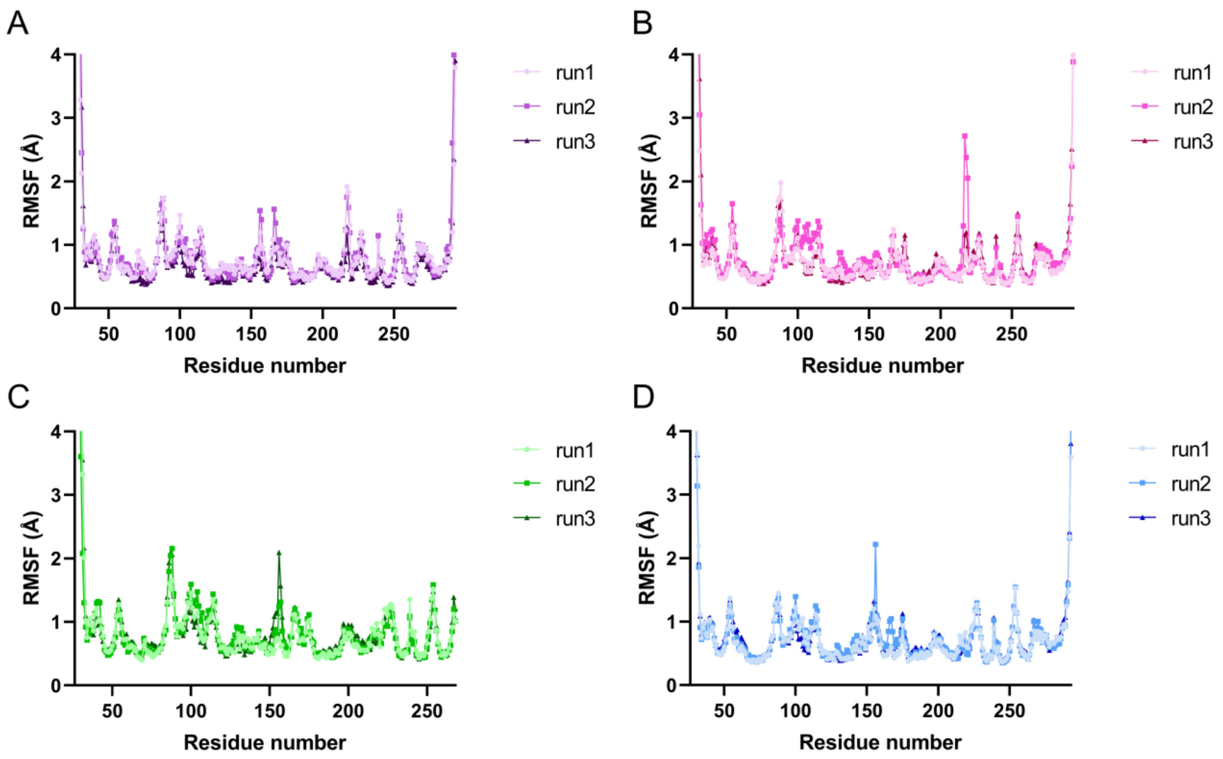


**Figure S8. Per residue RMSF plots for the KPC-2:meropenem-derived acyl-enzyme complex over triplicate 500 ns simulations. A.** KPC-2$^{E166Q}$:meropenem-derived acyl-enzyme (Δ2 tautomer) **B.** KPC-2:meropenem-derived acyl-enzyme (Δ2 tautomer) **C.** KPC-2$^{E166Q}$:meropenem-derived acyl-enzyme (Δ1-(2*R*) tautomer) **D.** KPC-2:meropenem-derived acyl-enzyme (Δ1-(2*R*) tautomer).

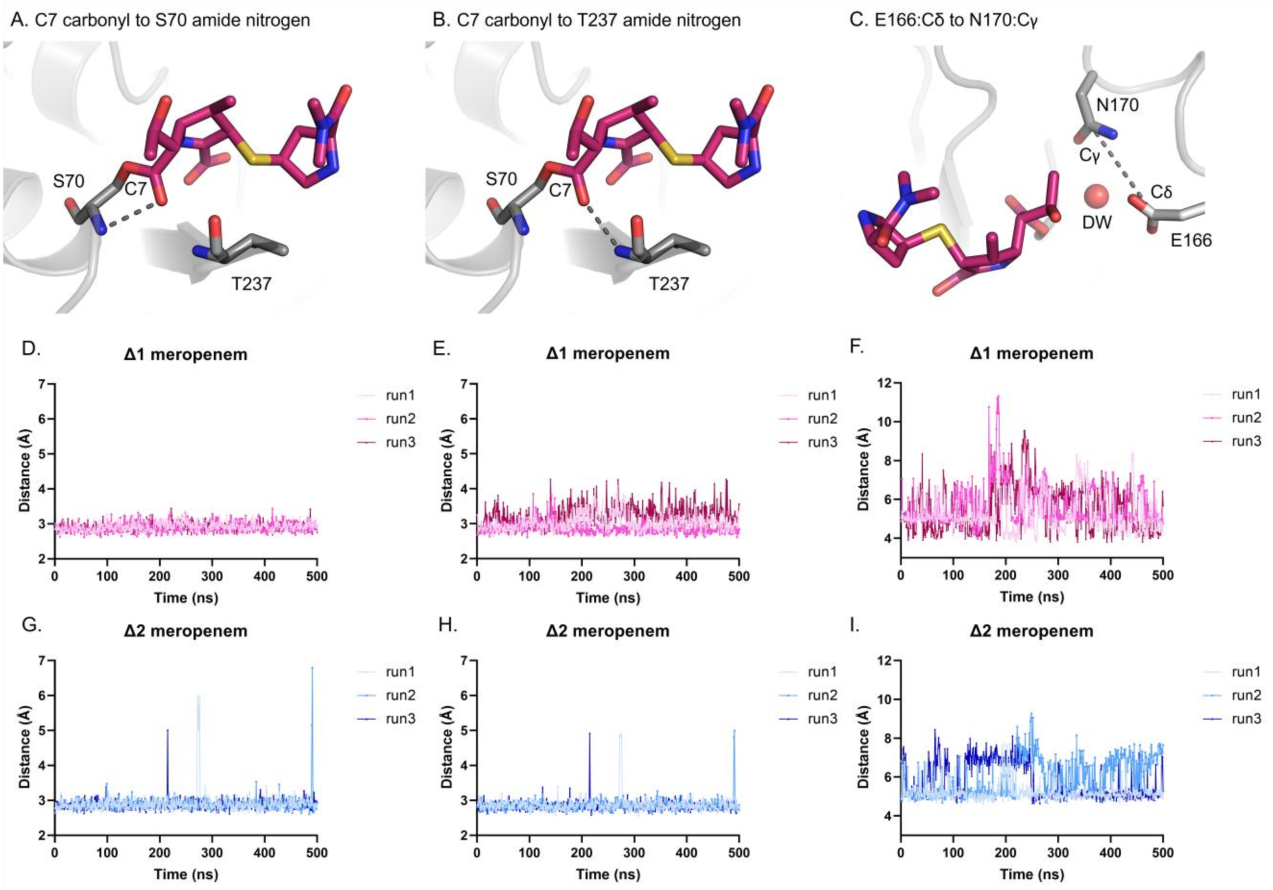


**Figure S9. Distance analyses of interactions of KPC-2:meropenem-derived acyl-enzymes in triplicate MM MD simulations. A. B** and **C,** starting structures of KPC-2 with the Δ1-(2*R*) meropenem-derived complex show the distances analyzed over the 500 ns trajectories. Relevant atoms and residues are labelled and the distances are represented as grey dashes. The meropenem-derived acyl-enzyme is shown as pink sticks and the KPC-2 protein as grey sticks and cartoon. The deacylating water (DW) is represented by a red sphere. **A, D** and **G,** Ser70 nitrogen (in the oxyanion hole) to the meropenem-derived C7 carbonyl distance. **B, E** and **H,** Thr237 nitrogen (oxyanion hole) to the meropenem-derived C7 carbonyl. **C, F** and **I,** distance between Cδ of Glu166 and Cγ of Asn170. **D, E** and **F,** respective distances over 500 ns trajectories between atoms in the Δ1-(2*R*) meropenem-derived acyl-enzyme shown in A, B and C. **G, H** and **I**: respective distances over 500 ns trajectories between atoms in the Δ2 meropenem-derived acyl-enzyme shown in A, B and C.

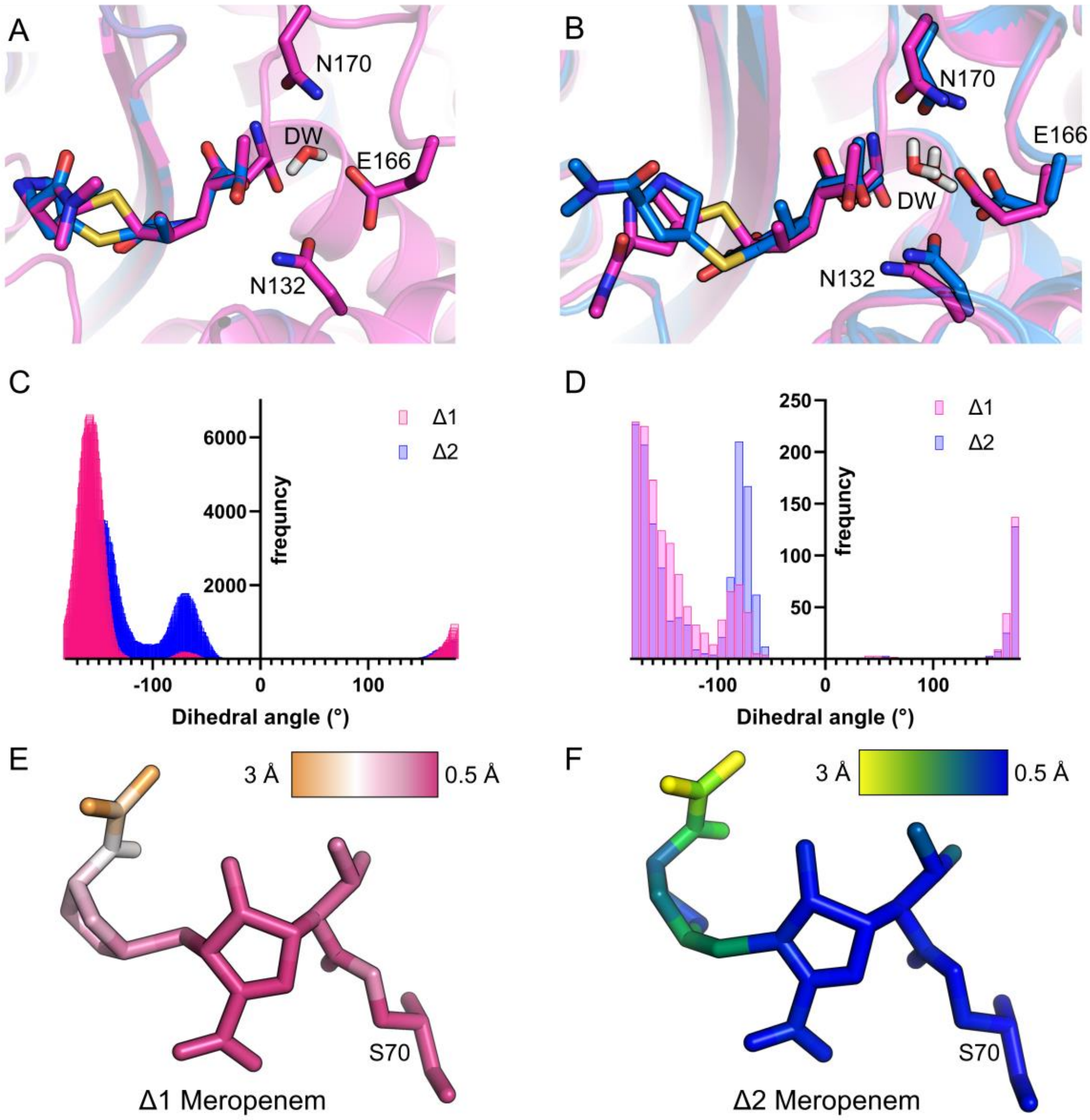


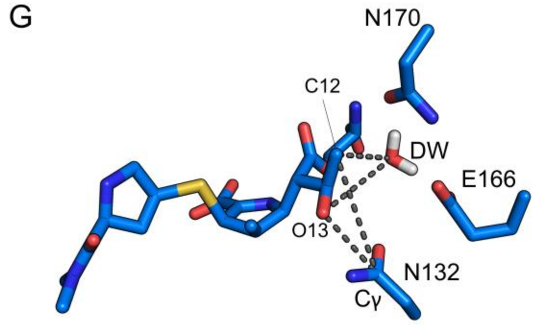


H

| | Δ1 | Δ2 |
|---|---|---|
| **N132 Cγ – Mer C12** | 5.15 (0.44) | 4.79 (0.53) |
| **N132 Cγ – Mer O13** | 3.88 (0.46) | 3.67 (0.25) |
| **DW – Mer C12** | 3.25 (0.38) | 3.54 (0.55) |
| **DW – Mer O13** | 4.36 (0.38) | 3.96 (0.49) |

**Figure S10. Position of the hydroxyethyl in MM and QM simulations. A.** Minimized starting structures. **B.** Equilibrated starting structures (following 1 ns MM MD). Histogram of the hydroxyethyl dihedral angle in **C.** Triplicate Adaptive string method (ASM) simulations and **D**. Triplicate molecular dynamics simulations. The Y axis represents the frequency (in number of frames). For ASM this included all sampled frames, for MD this included a frame for every ns of simulation. **E.** Ligand RMSF in ASM calculations shown on Δ1-(2*R*) meropenem (color ramped from orange to pink). **F**. Ligand RMSF in ASM calculations shown on Δ2 meropenem (color ramped from yellow to blue). **G.** Pymol representation of the distances (shown as dashes on the KPC-2:Δ2meropenem structure) measured between the 6α-hydroxyethyl of meropenem to the Deacylating water (DW) and Asn132 in ASM simulations. **H**. Corresponding distances in Å represented in **panel G**.

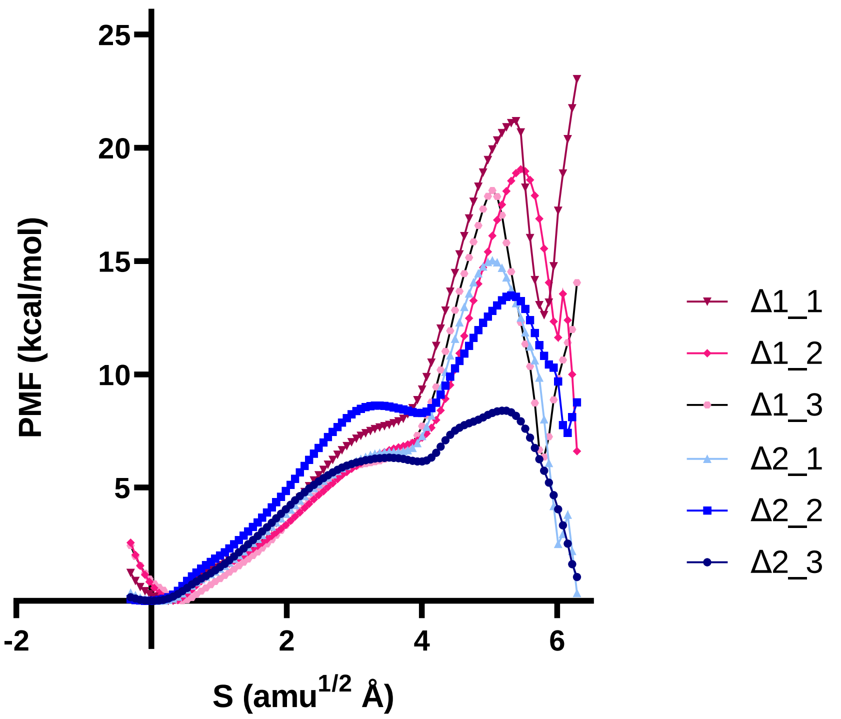


**Figure S11. ASM calculations for tetrahedral intermediate formation from the meropenem-derived acyl-enzyme in KPC-2 catalysis.** Potential of mean force (PMF) calculated by the ASM in kcal/mol is measured along the MFEP trajectory. Triplicate runs of each tautomer are coloured as pink (Δ1-(2*R*)) and blue (Δ2). The transition state (TS) is described as the high energy saddle point through the MFEP calculation before formation of the tetrahedral intermediate (TI).

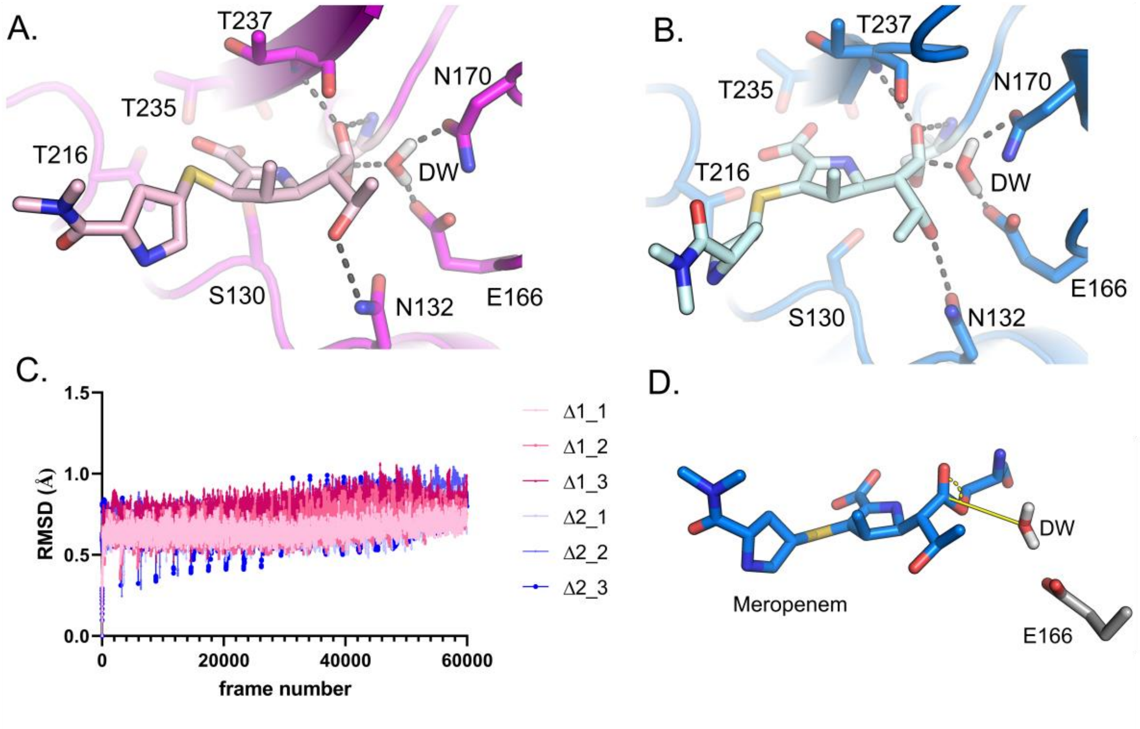


**Figure S12. Transition state analysis of adaptive string method calculations.** Snapshots of transition states with hydrogen bonding networks involving N132, E166, N170, T237 and the deacylating water (DW) shown as grey dashes. **A.** Δ1-(2*R*) meropenem-derived acyl-enzyme and **B.** Δ2 meropenem-derived acyl-enzyme**. C.** Acyl-enzyme RMSD during the adaptive string method (ASM) triplicate trajectories. **D.** The deacylating water (DW) to C7 carbonyl angle was measured during the ASM simulations. The average DW-C7 angle sampled in Δ1-(2*R*) meropenem simulations was 112.5°±3.4 and for Δ2 111.7°±3.6, yet Δ2 sampled the favored 107°-109° Burgi-Dunitz angle in 3.2% more frames (Δ2= 23.8% vs Δ1=20.6%) over the ASM trajectories.

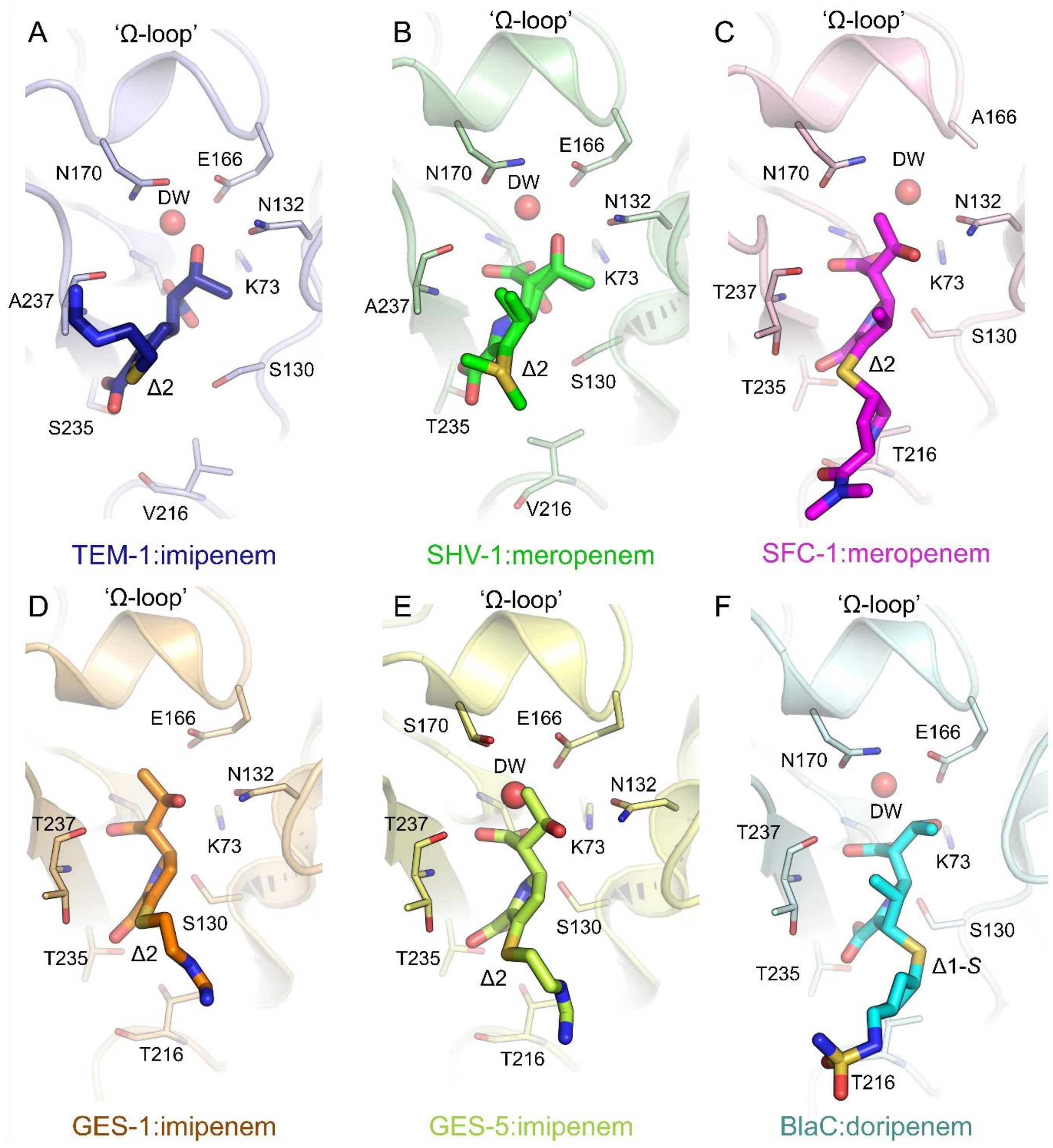


**Figure S13. Comparisons of the interactions of carbapenem acyl-enzyme complexes within Class A SBLs. A.** TEM-1:imipenem-derived acyl-enzyme complex (PDB:1BT5)[2], blue. **B.** SHV-1:meropenem-derived acyl-enzyme complex (PDB:2ZD8)[3], green. **C.** SFC-1:meropenem-derived acyl-enzyme complex, pink (featuring A166 mutation PDB:4EV4))[4]. **D.** GES-1:imipenem-derived acyl-enzyme complex (PDB:4GOG)[5], orange. **E.** GES-5:imipenem-derived acyl-enzyme complex (PDB:4H8R)[5], lime. **F.** BlaC:doripenem-derived acyl-enzyme complex (PDB:3IQA)[6], cyan. Carbapenem-derived acyl-enzymes are highlighted as bold sticks and key active site amino acids (if present in the structure) are labelled. The tautomeric state (Δ1-(2*R*) or Δ2) of the carbapenem-derived acyl-enzyme is labelled adjacent to the C2-sulphur. Waters closest to the putative deacylating position (if present) are shown as red spheres and labelled DW.

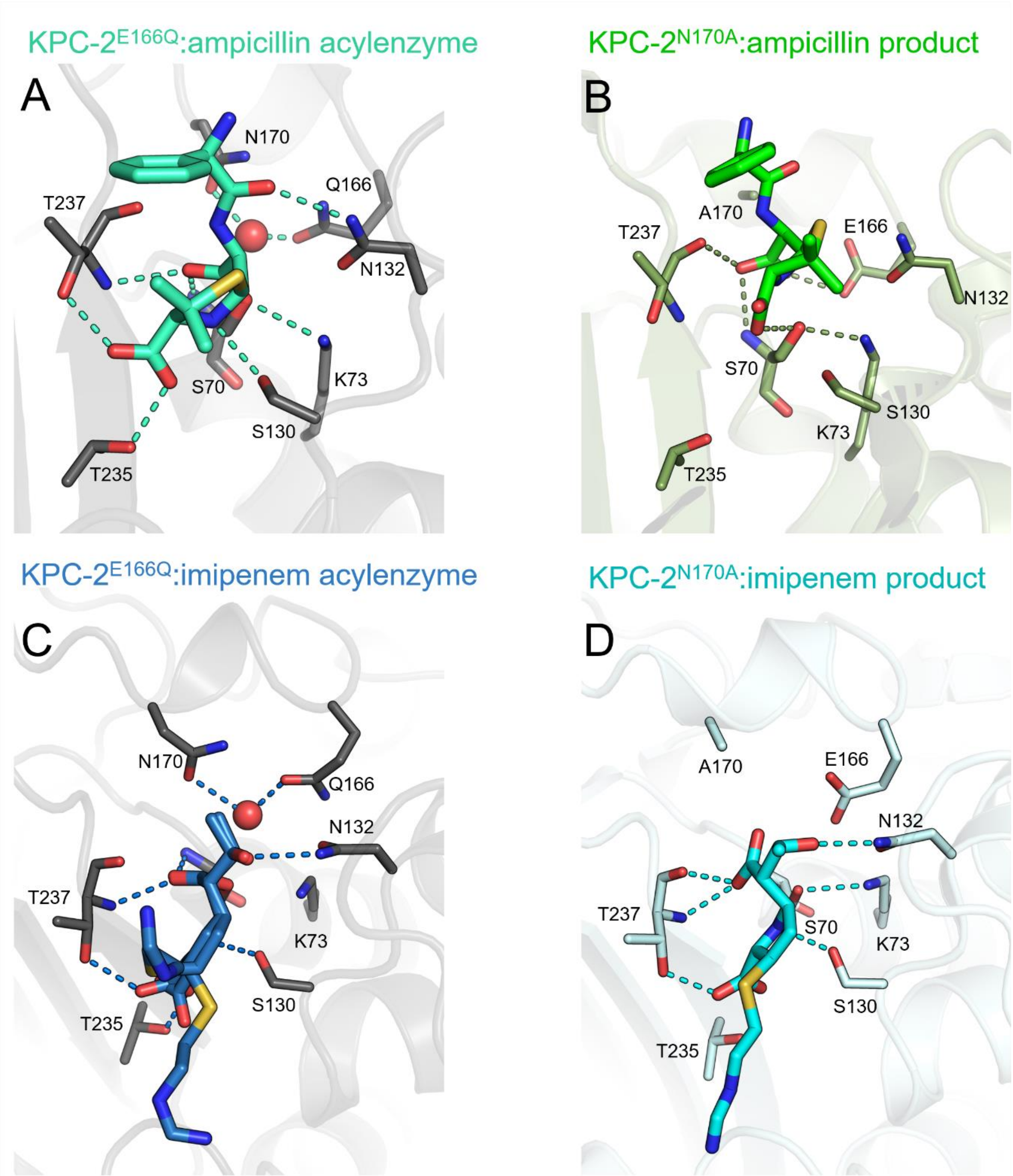


**Figure S14. Comparison of KPC-2:antibiotic acyl-enzyme and non-covalent hydrolysis product complexes.** Antibiotics and important active site amino acids are highlighted as sticks, possible H-bonds are identified as dashes. **A.** KPC-2$^{E166Q}$:ampicillin acyl-enzyme complex (teal). **B.** KPC-2$^{N170A}$:ampicillin non-covalent hydrolysis product (green, PDB:6XD7)[7]. **C.** KPC-2$^{E166Q}$:imipenem acyl-enzyme complex (blue). Note the two conformations of the carboxylate in this case. **D.** KPC-2$^{N170A}$:imipenem non covalent hydrolysis product (cyan, PDB 6XJ8)[7]. Note, mutation of Asn170 to Ala results in loss of a water in the deacylating position.

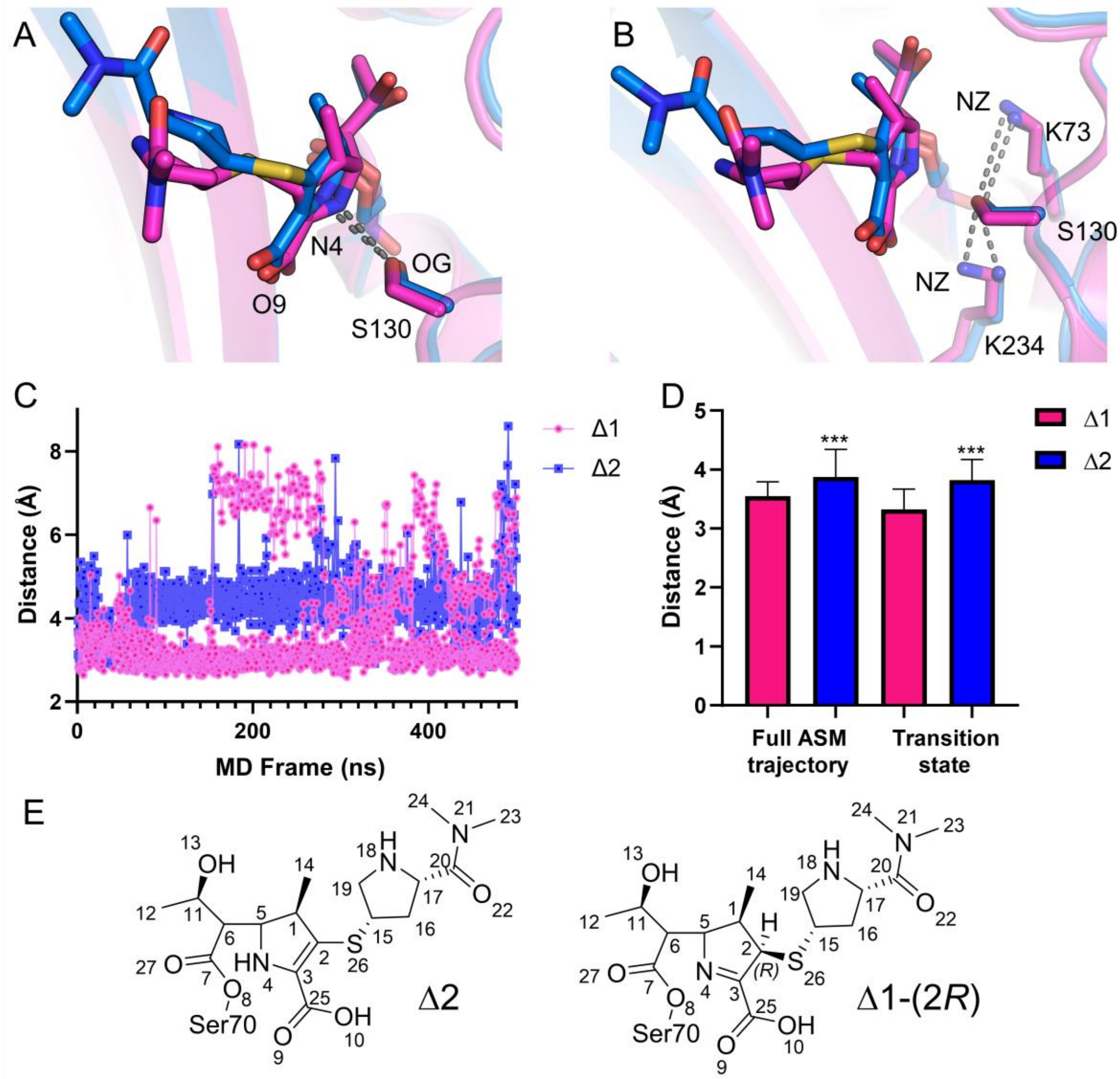


F

| Acceptor | Donor | Δ1 Full Traj % | Δ2 Full Traj % | Δ1 TS % | Δ2 TS % |
|---|---|---|---|---|---|
| Ser130 OG | Lys234 NZ | 56.09 | 26.92 | 42.99 | 11.55 |
| Ser130 OG | Lys73 NZ | 7.77 | 10.5 | 10.46 | 9.52 |
| Mero N4 | Ser130 OG | 2.27 | 0.09 | 6.88 | 0.1 |
| Mero O9 | Ser130 OG | 95.95 | - | 87.36 | - |

**Figure S15. Ser130 interactions throughout MD and ASM simulations**. A and B. Equilibrated starting structures (following 1 ns MM MD) Δ1-(2*R*), pink and Δ2, blue showing S130 OG interactions with N4 of meropenem and NZ of Lys73 and Lys 234 respectively. C. Distance analysis of S130 to N4 in triplicate 500ns MD simulations, average distances are Δ1-(2*R*) =3.76 (±1.27) Å and Δ2=4.38 (0.59). D. Distance between meropenem N-4 and Ser130-OG in ASM simulations. Significance determined with unpaired t-test performed in GraphPad Prism**,** a P value of < 0.0001. E. Δ1-(2*R*) and Δ2 acyl-enzyme representations with numbered atoms corresponding to panel F. F. Hydrogen bonding interactions with Ser130 during ASM simulations for the full trajectory compared to the transition state.

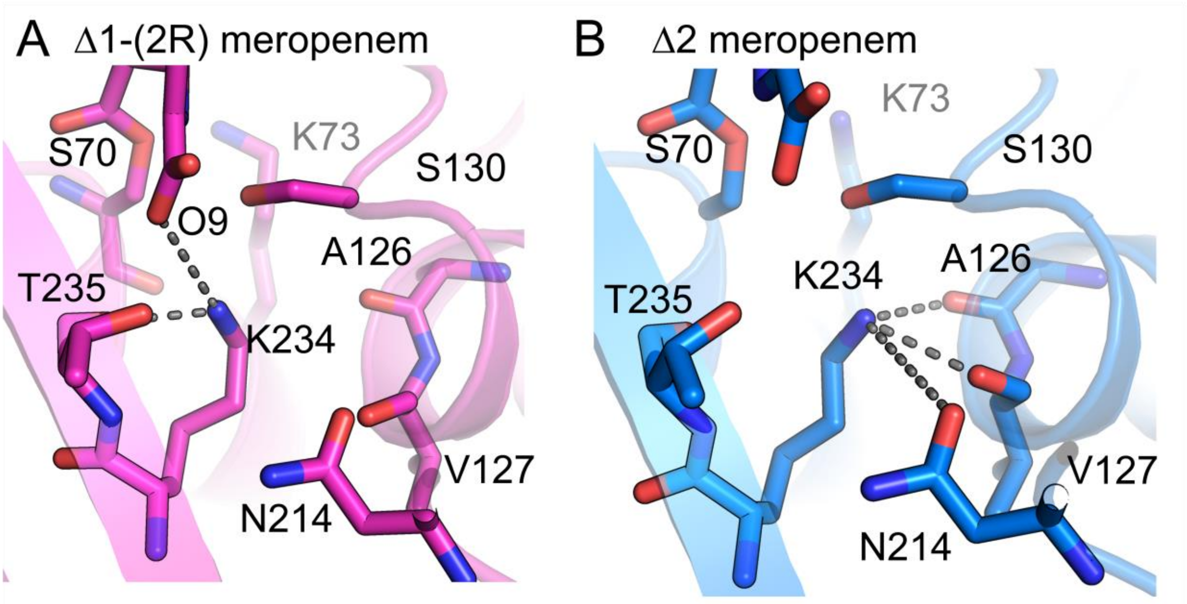


C

| Acceptor | Donor | Δ1 Full Traj % | Δ2 Full Traj % | Δ1 TS % | Δ2 TS % |
|---|---|---|---|---|---|
| Val127 O | Lys234 NZ | 0 | 74.26 | 0 | 66.85 |
| Asn214 OD1 | Lys234 NZ | 0 | 16.4 | 0 | 0 |
| Ala126 O | Lys234 NZ | 0 | 15.56 | 0 | 31.37 |
| Thr235 O | Lys234 NZ | 79.75 | 0 | 85.55 | 0 |
| Mer O9 | Lys234 NZ | 5.27 | 0 | 6.74 | 0 |

**Figure S16. Lys234 networks throughout ASM simulations.** Representations of the distinct hydrogen bonding networks to K234 in ASM simulations **A**. Δ1-(2*R*) meropenem acyl-enzyme (equilibrated starting structure) and **B**. Δ2 meropenem acyl-enzyme (equilibrated starting structure). **C**. Distinct hydrogen bonding networks from K234, represented by % of frames.

**Supporting note 1**

Four systems (KPC-$2^{E166Q}$:meropenem-Δ2, KPC-2:meropenem-Δ2, KPC-$2^{E166Q}$:meropenem- Δ1-(2*R*) and KPC-2:meropenem- Δ1-(2*R*)) were energy minimised and equilibrated prior to 500 ns triplicate simulations (1.5 µs total) and analysed in cpptraj. Backbone ($C_{\alpha}$) root mean square deviation (RMSD) values of residues 25-291 remained below 1.7 Å throughout 500 ns trajectories (**Figure S6**). The N- and C-termini displayed significant mobility in RMSD and root mean square fluctuation (RMSF) calculations (**Figures S6 and S7**) consistent with ours and others' apoenzyme simulations and analyses of KPC-2[1-2]. Small differences in RMSF around residues 216 and 217 (the 'hinge region') were visible between simulations of the Δ1-(2*R*) and Δ2 tautomer (**Figures 5 and S7**) which upon visual inspection appeared mobile throughout the duration of MD simulations[2]. Ω-Loop RMSF remained below 1.5 Å and stable in the presence of meropenem, unlike that observed for the 'bulky' oxyimino cephalosporin ceftazidime.[2] This is consistent with the experimental electron density for this region within the corresponding crystal structures (**Table S1**).

Distance measurements between backbone amide atoms (Ser70:N and Thr237:N) and the C7 carbonyl of the meropenem-derived acyl-enzyme in both Δ1-(2*R*) and Δ2 tautomers over the triplicate 500 ns MD simulations revealed these distances (and subsequent likely hydrogen bonding interactions) remained relatively stable at around 3 Å (**Figure S8**). With the Δ2 meropenem-derived acyl-enzyme, the C7 carbonyl lost interaction with the oxyanion hole in 2 frames, representing a rotation of this region in a similar orientation to that seen in crystal structures of SHV-1 and TEM-1.[3-4] This movement may also link to recent studies of KPC-2 that indicated permissive and non-permissive states may be possible during carbapenem hydrolysis.[5] Extensive analysis of Molecular Dynamics trajectories reveals no large differences in behaviour of the KPC-2:meropenem acyl-enzyme between the Δ1-(2*R*) and Δ2 tautomers. These results were expected due to the minute differences in protonation state of C3 and N4 of the carbapenem-derived scaffold (**Figure 3**).

Higher level computational approaches (combined Quantum mechanics/ Molecular mechanics, QM/MM) were therefore utilised to elucidate differences which may exist between the tautomers of the meropenem-derived acyl-enzymes. Hybrid QM/MM umbrella sampling simulations were used to investigate meropenem hydrolysis in the KPC-2 acyl-enzyme.

In a protocol that we have previously shown to be effective for modeling deacylation of carbapenem-derived acyl-enzymes by class A β-lactamases[12], we employ the SCC-DFTB2[13-15] QM method in the AMBER16 simulation package,[16] using the ff14SB[17] MM forcefield for protein, the TIP3P-Ew[18] water model and the General AMBER Force Field (GAFF)[16] for meropenem-derived regions not in the QM region. The QM region (modelled with a charge of -2) comprises the water in the deacylating position (DW), Ser70, Glu166 and the common carbapenem scaffold. Conventional QM/MM umbrella sampling MD was performed along two distance-based reaction coordinates to simulate the deacylation reaction (**Figure S5**). The first is the linear combination of distances between the distance of Glu166 OE2 and the nearest proton of the deacylating water and the distance between the same proton and the oxygen of the deacylating water, 1: rx = d(OεGlu166–HDW) – d(HDW–ODW) which corresponds to proton transfer from DW to Glu166. The second is the distance from the newly formed hydroxide to the C8 carbonyl, 2: = d(C7meropenem–ODW) which represents the nucleophilic attack of DW on the carbonyl carbon (**Figure S5**). 20 ps of QM/MM MD were performed for each simulation window, and simulations repeated in triplicate. 2D minimum free energy paths (MFEPs) were calculated as previously described[12], using the weighted-histogram analysis method (WHAM); the highest point (saddle point) along the MFEP is taken as the transition state, giving the activation free energy, $\Delta^{\ddagger}G_{\text{calc}}$ **Table S4**). The adaptive string method (ASM) path calculation method was also used for MFEP calculation of the first step of carbapenem deacylation. Two collective variables were used with the ASM, with the first collective variable subtly differing to the first reaction coordinate used for conventional US: The distance between Glu166 OE2 and a proton of the deacylating water, 1: rx = d(OεGlu166–HDW) and the distance between the meropenem C7 and the oxygen of the newly formed hydroxide, 2: ry = d(C7meropenem–ODW). Each node along the string was sampled for 60 ps and the simulations repeated in triplicate.

**Supporting note 2**

We measured the Ser130-OG to N4 distance over ASM trajectories derived from QM/MM MD simulations of the respective acyl-enzymes, finding significant differences both across the whole trajectory and between the respective deacylation transition states (**Figure S15**). Analysis of H-bond networks determined that Ser130 is likely positioned, by Lys73 and Lys234, to interact with either the meropenem C3 carboxylate or N4 over both the whole trajectory and the transition state frames. Indeed, further analysis of the orientation of Ser130 throughout the ASM trajectories also reveals more consistent hydrogen bonding to the Lys234 side chain amide (Lys234-NZ) in the Δ1-(2*R*) (imine) tautomer than in the Δ2 (enamine) (**Figure S15**). Thus, more stable electrostatic interactions in the Δ1-(2*R*) acyl-enzyme sustain Ser130-OG in a position close to either meropenem-O9 or meropenem N-4, whilst the relative instability of the Δ2 acyl-enzyme allows the Ser130 side chain to explore conformational space further from meropenem N-4. These differences directly result from differences between tautomers in the hydrogen bonding network around the Lys234 side chain (**Figure S16**). Specifically, Lys234-NZ makes interactions exclusive to each tautomer; hydrogen bonding predominantly to the Thr235 backbone oxygen in the Δ1-(2*R*), and to the Val127 backbone oxygen in the Δ2 configuration (**Figure S16**). In consequence different orientations of the Lys234 side chain prevail in each tautomer, subsequently hindering its propensity to hydrogen bond to Ser130-OG in Δ2 but promoting it in Δ1-(2*R*), accounting for the difference between tautomers in the distance between Ser130-OG and meropenem N-4. Alongside the H-bond networks around T216, T235 (as described in results) our simulations reveal that the Δ1-(2*R*) meropenem acyl-enzyme makes more extensive hydrogen bonding interactions with KPC-2, and is consequently more stable, than the Δ2, resulting in an increased free energy barrier to tetrahedral intermediate formation, as evidenced by the 7.1 kcal/mol difference between tautomers identified by ASM calculations.

**Supporting Information References**


1. Pemberton, O. A.; Zhang, X.; Chen, Y., Molecular Basis of Substrate Recognition and Product Release by the Klebsiella pneumoniae Carbapenemase (KPC-2). *J Med Chem* **2017,** *60* (8), 3525-3530.
2. Maveyraud, L.; Mourey, L.; Kotra, L. P.; Pedelacq, J.-D.; Guillet, V.; Mobashery, S.; Samama, J.-P., Structural Basis for Clinical Longevity of Carbapenem Antibiotics in the Face of Challenge by the Common Class A β-Lactamases from the Antibiotic-Resistant Bacteria. *Journal of the American Chemical Society* **1998,** *120* (38), 9748-9752.
3. Nukaga, M.; Bethel, C. R.; Thomson, J. M.; Hujer, A. M.; Distler, A.; Anderson, V. E.; Knox, J. R.; Bonomo, R. A., Inhibition of class A beta-lactamases by carbapenems: crystallographic observation of two conformations of meropenem in SHV-1. *J Am Chem Soc* **2008,** *130* (38), 12656-62.
4. Fonseca, F.; Chudyk, E. I.; van der Kamp, M. W.; Correia, A.; Mulholland, A. J.; Spencer, J., The Basis for Carbapenem Hydrolysis by Class A β-Lactamases: A Combined Investigation using Crystallography and Simulations. *Journal of the American Chemical Society* **2012,** *134* (44), 18275-18285.
5. Smith, C. A.; Frase, H.; Toth, M.; Kumarasiri, M.; Wiafe, K.; Munoz, J.; Mobashery, S.; Vakulenko, S. B., Structural Basis for Progression toward the Carbapenemase Activity in the GES Family of β-Lactamases. *Journal of the American Chemical Society* **2012,** *134* (48), 19512-19515.
6. Tremblay, L. W.; Fan, F.; Blanchard, J. S., Biochemical and structural characterization of Mycobacterium tuberculosis beta-lactamase with the carbapenems ertapenem and doripenem. *Biochemistry* **2010,** *49* (17), 3766-73.
7. Mehta, S. C.; Furey, I. M.; Pemberton, O. A.; Boragine, D. M.; Chen, Y.; Palzkill, T., KPC-2 β-lactamase enables carbapenem antibiotic resistance through fast deacylation of the covalent intermediate. *The Journal of biological chemistry* **2021,** *296*, 100155.
8. Galdadas, I.; Qu, S.; Oliveira, A. S. F.; Olehnovics, E.; Mack, A. R.; Mojica, M. F.; Agarwal, P. K.; Tooke, C. L.; Gervasio, F. L.; Spencer, J.; Bonomo, R. A.; Mulholland, A. J.; Haider, S., Allosteric communication in class A β-lactamases occurs via cooperative coupling of loop dynamics. *eLife* **2021,** *10*, e66567.
9. Tooke, C. L.; Hinchliffe, P.; Bonomo, R. A.; Schofield, C. J.; Mulholland, A. J.; Spencer, J., Natural variants modify Klebsiella pneumoniae carbapenemase (KPC) acyl-enzyme conformational dynamics to extend antibiotic resistance. *The Journal of biological chemistry* **2021,** *296*, 100126.
10. Nukaga, M.; Bethel, C. R.; Thomson, J. M.; Hujer, A. M.; Distler, A.; Anderson, V. E.; Knox, J. R.; Bonomo, R. A., Inhibition of Class A β-Lactamases by Carbapenems: Crystallographic Observation of Two Conformations of Meropenem in SHV-1. *Journal of the American Chemical Society* **2008,** *130* (38), 12656-12662.
11. Cortina, G. A.; Hays, J. M.; Kasson, P. M., Conformational Intermediate That Controls KPC-2 Catalysis and Beta-Lactam Drug Resistance. *ACS Catalysis* **2018,** *8* (4), 2741-2747.
12. Chudyk, E. I.; Limb, M. A. L.; Jones, C.; Spencer, J.; van der Kamp, M. W.; Mulholland, A. J., QM/MM simulations as an assay for carbapenemase activity in class A β-lactamases. *Chemical Communications* **2014,** *50* (94), 14736-14739.
13. de M Seabra, G.; Walker, R. C.; Elstner, M.; Case, D. A.; Roitberg, A. E., Implementation of the SCC-DFTB method for hybrid QM/MM simulations within the amber molecular dynamics package. *J Phys Chem A* **2007,** *111* (26), 5655-64.
14. Woodcock, H. L.; Hodoscek, M.; Brooks, B. R., Exploring SCC-DFTB paths for mapping QM/MM reaction mechanisms. *J Phys Chem A* **2007,** *111* (26), 5720-8.
15. Elstner, M.; Seifert, G., Density functional tight binding. **2014,** *372* (2011), 20120483.
16. D.A. Case, J. T. B., R.M. Betz, D.S. Cerutti, T.E. Cheatham, III, T.A. Darden, R.E. Duke, T.J. Giese, H. Gohlke, A.W. Goetz, N. Homeyer, S. Izadi, P. Janowski, J. Kaus, A. Kovalenko, T.S. Lee, S. LeGrand, P. Li, T. Luchko, R. Luo, B. Madej, K.M. Merz, G. Monard, P. Needham, H. Nguyen, H.T. Nguyen, I. Omelyan, A. Onufriev, D.R. Roe, A. Roitberg, R. Salomon-Ferrer, C.L. Simmerling, W. Smith, J. Swails, R.C. Walker, J. Wang, R.M. Wolf, X. Wu, D.M. York and P.A. Kollman, AMBER 2015. University of California, San francisco, 2015.

17. Maier, J. A.; Martinez, C.; Kasavajhala, K.; Wickstrom, L.; Hauser, K. E.; Simmerling, C., ff14SB: Improving the Accuracy of Protein Side Chain and Backbone Parameters from ff99SB. *J Chem Theory Comput* **2015,** *11* (8), 3696-713.
18. Jorgensen, W. L.; Chandrasekhar, J.; Madura, J. D.; Impey, R. W.; Klein, M. L., Comparison of simple potential functions for simulating liquid water. *The Journal of Chemical Physics* **1983,** *79* (2), 926-935.
